# Endogenous RNAi silences a burgeoning sex chromosome arms race

**DOI:** 10.1101/2022.08.22.504821

**Authors:** Jeffrey Vedanayagam, Ching-Jung Lin, Ranjith Papareddy, Michael Nodine, Alex S. Flynt, Jiayu Wen, Eric C. Lai

**Affiliations:** Developmental Biology Program, Sloan Kettering Institute, 430 East 67th St, ROC-10, New York, NY 10065, USA; Weill Graduate School of Medical Sciences, Weill Cornell Medical College, New York, New York 10065, USA; Gregor Mendel Institute (GMI), Austrian Academy of Sciences, Vienna Biocenter (VBC), 1030 Vienna, Austria; Department of Biological Sciences University of Southern Mississippi Hattiesburg, Mississippi 39406, USA; Department of Genome Sciences, The John Curtin School of Medical Research, The Australian National University, Canberra, Australia

## Abstract

Although the biological utilities of endogenous RNAi (endo-RNAi) have been largely elusive, recent studies reveal its critical role in the non-model fruitfly *Drosophila simulans* to suppress selfish genes, whose unchecked activities can severely impair spermatogenesis. In particular, hairpin RNA (hpRNA) loci generate endo-siRNAs that suppress evolutionary novel, X-linked, meiotic drive loci. The consequences of deleting even a single hpRNA (*Nmy*) in males are profound, as such individuals are nearly incapable of siring male progeny. Here, comparative genomic analyses of *D. simulans* and *D. melanogaster* mutants of the core RNAi factor *dcr-2* reveal a substantially expanded network of recently-emerged hpRNA-target interactions in the former species. The *de novo* hpRNA regulatory network in *D. simulans* bears compelling signatures of sex chromosome conflict and provides insight into molecular strategies that underlie hpRNA emergence. In particular, our data support the existence of ongoing rapid evolution of Nmy/Dox-related networks, recurrent targeting of testis HMG Box loci by hpRNAs, and connections to the piRNA pathway. Importantly, the impact of the endo-RNAi network on gene expression flips the convention for regulatory networks, since we observe strong derepression of targets of the youngest hpRNAs, but only mild effects on the targets of the oldest hpRNAs. These data suggest that endo-RNAi are especially critical during incipient stages of intrinsic sex chromosome conflicts, and that continual cycles of distortion and resolution may contribute to the segregation of species.

## Introduction

In sexually reproducing organisms, structurally distinct sex chromosomes (X/Y or Z/W) are involved in sex-specific regulatory processes, such as sex determination and dosage compensation (Bachtrog, 2020; Charlesworth, 1991). The genomic distinction of sex chromosomes, compared to their autosomal counterparts, underlies strikingly contrasting features including (1) reduction or lack of recombination, (2) strategies to equalize gene expression from the X or Z of males and females and (3) accumulation of repeats on the degenerating Y (or W) chromosome (Abbott et al., 2017). Accordingly, XY and ZW chromosomes are especially evolutionarily dynamic (Johnson and Lachance, 2012).

The rapid and continual evolution and emergence of sex chromosomes, along with their contrasting biological interests and fates, is linked to their involvement in intragenomic conflict (Meiklejohn and Tao, 2010) and sex chromosome meiotic drive (Bachtrog, 2020). In particular, selfish sex-linked genes can impair transmission of the reciprocal sex chromosome, thereby favoring the driving chromosome amongst progeny. Sex chromosome meiotic drive can be easily observed in deviation of *sex-ratio* (SR) from equality (Jaenike, 2008). Fisher’s principle proposes that, if males and females cost equal amounts to produce, an equal ratio of the sexes will be the equilibrium (Fisher, 1930). However, SR drive systems have been widely documented, indicating recurrence of sex chromosome drive in nature. Fisher proposed that sex-biased populations direct their reproductive efforts disproportionately to the rarer sex, thus tending towards normalization of SR over the long term. However, the molecular bases of SR distortion and restoration of parity are poorly understood. This is due in part due to the fact that, despite their ubiquity in nature, many well-studied model organisms lack documented SR drive systems. For example, even though mutants of sex determination or dosage compensation systems can distort the sex of viable progeny, extensive studies have not uncovered strong, selfish SR drive loci in the well-studied fruitfly *D. melanogaster* (*Dmel*).

Curiously then, a history of genetic analyses uncovered three independent sex chromosome drive systems in *D. simulans* (*Dsim*), a close sister species of *Dmel* (Presgraves and Meiklejohn, 2021). *Dsim* bears the Winters, Durham, and Paris systems, meiotic drive systems that map to distinct genomic intervals and indicate multiple newly-emerged strategies that deplete male progeny (Cazemajor et al., 2000; Tao et al., 2007a; Tao et al., 2001; Tao et al., 2007b). Despite progress on the identification of potential drivers and/or suppressors for the three SR drive systems (Helleu et al., 2016; Lin et al., 2018; Muirhead and Presgraves, 2021; Tao *et al*., 2007a; Tao *et al*., 2007b; Vedanayagam et al., 2021), much remains to be understood regarding their molecular mechanisms, and even whether SR meiotic drive loci have been comprehensively identified in this species. The limited genetic tools and genomic data in this non-model fruitfly have impeded efforts, even though *Dsim* is arguably the premier model to explore the molecular bases of SR drive.

Recently, we revealed that two genetically identified loci that suppress SR drive encode hairpin RNA (hpRNAs), which generate endogenous siRNAs (endo-siRNAs). In particular, the Winters SR suppressor (*Nmy*) and the Durham SR suppressor (*Tmy*) encode related hpRNAs that have capacities to silence the SR distorter *Dox* and its paralog *MDox* (Lin *et al*., 2018), to equalize SR in *Dsim*. Notably, both *Dox* and *MDox* are located on the X chromosome and are silenced by endo-siRNAs in *Dsim* testis. These attributes fit the proposition that meiotic drivers may preferentially be encoded on the X, and exploit male meiosis to gain unfair transmission advantages. Moreover, the family of Dox-related genes and related hpRNAs has undergone massive amplification in the *simulans*-clade sister species *D. sechellia* (*Dsech*) and *D. mauritiana* (*Dmau*), but none of these driver or suppressor loci are present in the *Dmel* genome (Muirhead and Presgraves, 2021; Vedanayagam *et al*., 2021). These findings are testament to the rapid evolution (both emergence and disappearance) of SR meiotic drive systems, and a key role for endo-RNAi to suppress incipient selfish genes located on the X.

To test for broader roles of RNAi in suppressing meiotic drive and/or sex chromosome conflict, we used short and long transcriptome data from mutants of the core RNAi factor *Dcr-2* to perform a functional evolutionary comparison of hpRNA regulatory networks in *Dmel* and *Dsim* testis. We reveal asymmetric proliferation of evolutionarily novel hpRNAs in *Dsim*, which preferentially repress *de novo* X-linked genes, indicating a burgeoning sex chromosome conflict in this species. These loci provide insights into the earliest stages of molecular emergence of hpRNAs.

Surprisingly, the newest hpRNA-target interactions mediate much larger regulatory effects than the oldest hpRNA-target interactions, thereby inverting the convention of miRNA-mediated networks. Overall, we conclude that RNAi has a much larger role in silencing sex chromosome conflict than anticipated, and suggests that resolution of active intragenomic conflicts may contribute to speciation.

## Results

### Generation of *D. simulans dcr-2* deletion mutants marked by *white^+^*

We recently reported deletion alleles of core RNAi factors [*dcr-2* and *ago2*] in *D. simulans* (*Dsim*) (Lin *et al*., 2018). Although these mutants are viable, they are completely male sterile, and therefore cannot be maintained as stable stocks. This presents a technical challenge since *Dsim* lacks balancer chromosomes and the *3xP3:DsRed* marker was not fully reliable to distinguish heterozygotes from homozygotes. Therefore, preparation of pure homozygous material required extensive genotyping of small batches of dissected flies prior to combining samples for RNA isolation. Moreover, in initial RNA-seq analyses, genotyped samples were still prone to contamination. As a further complication, due to extremely high expression of many accessory gland transcripts (Brown et al., 2014), we noticed that even minute quantities of contaminating accessory gland could produce large biases in gene expression between libraries. Since *Dsim* mutants were generated in a *white* mutant background (*w[XD1*]), their testis was colorless and thus more challenging to visualize compared to a *white^+^* background, where the testis is bright yellow. For these reasons, the preparation of suitable quantities of *Dsim* RNAi mutant testis was not straightforward.

To address these issues, we used CRISPR/Cas9 to generate multiple founders of a new *Dsim Δdcr-2* null allele, where most of its coding region is replaced with *mini-white^+^* (*w^+^)* (**Supplementary Figure 1**). We anticipated selecting homozygotes with deeper eye color, as is typical in *Dmel*; however, this was also not fully reliable due to the red eyes of these alleles. Instead, by crossing *dcr-2* alleles marked by *3xP3:DsRed* and *w^+^*, we could select trans-heterozygotes carrying both dominant markers. Although DsRed^+^ eyes cannot be effectively scored in a *w^+^* background, it is still possible to score DsRed^+^ ocelli (**Supplementary Figure 1**). Our independent Δ*dcr-2[w^+^]* alleles were viable but specifically sterile in males, exhibited severe spermatogenesis defects, and failed to complement their corresponding DsRed alleles. We therefore used the trans-heterozygotes for subsequent analyses.

### Signature features of hpRNA loci in short/long RNAs from wildtype and *dcr-2* mutants

In contrast to *Dmel* RNAi mutants, which are viable and sub-fertile (Wen et al., 2015), deletion of core RNAi factors in *Dsim* result in complete male sterility (Lin *et al*., 2018). This is due at least in part to the requirements of *Nmy* and *Tmy*, which are *de novo* hpRNAs that silence incipient X chromosome *sex ratio* distorters (*Dox* and *MDox*) in the male germline (Lin *et al*., 2018). To assess the impact of RNAi loss more globally, and to compare *Dmel* and *Dsim* in greater detail, we generated biological replicates of small RNA and total RNA sequencing data from testis of *dcr-2* heterozygotes and mutants in *Dmel*, and *w[XD1]* vs. *dcr-2* mutants in *Dsim*. In control, we expect that primary hpRNA (pri-hpRNA) transcripts are cleaved by Dcr-2 into 21-22 nucleotide (nt) siRNAs, while *bona fide* siRNAs should be lost in *dcr-2* mutants concomitant with accumulation of their progenitor mRNAs (**Figure 1A** and **Supplementary Figure 2**).

**Figure 1.**
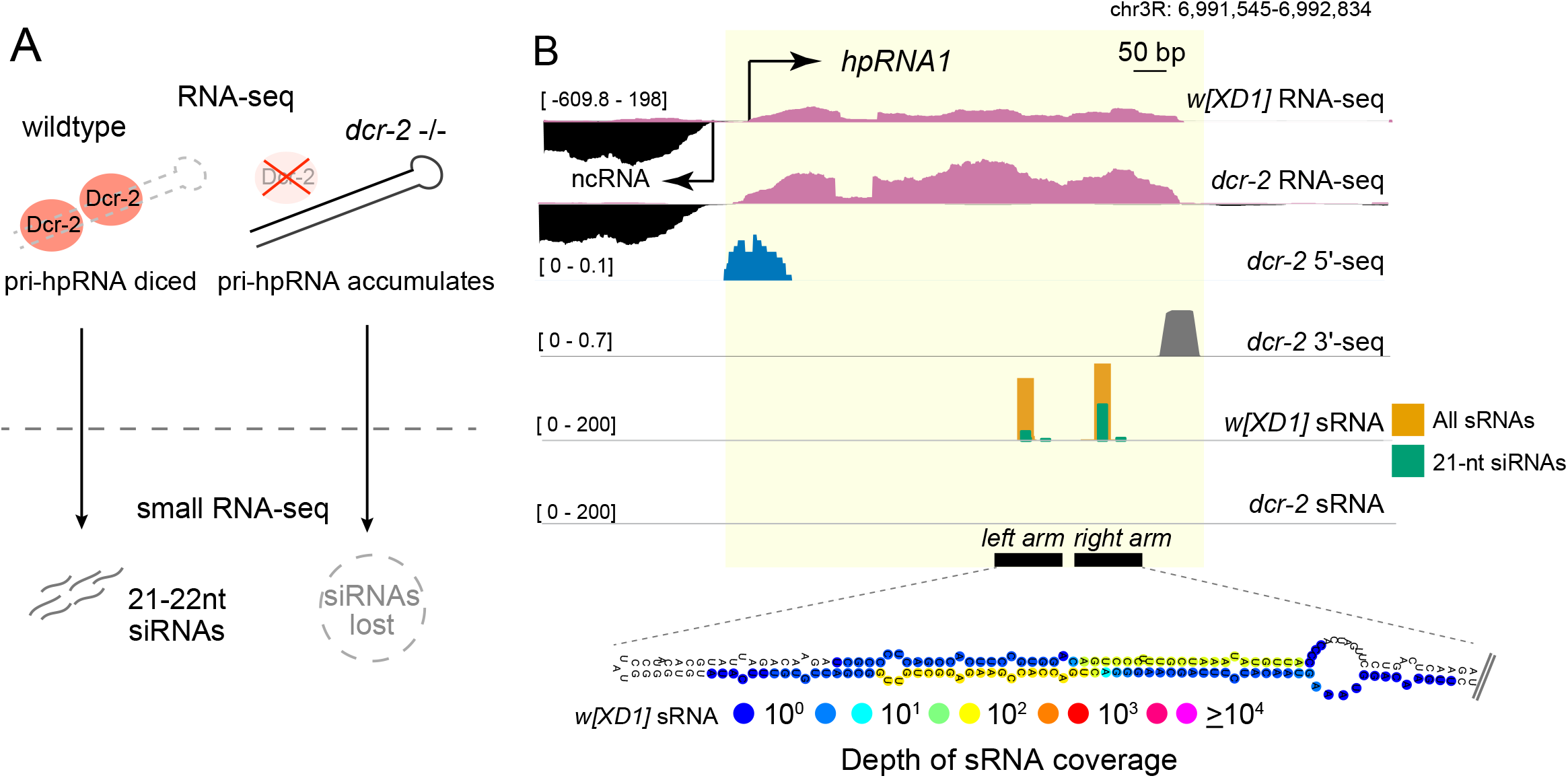
Annotation of novel hpRNAs using small RNA and RNA-seq data. (A) Schematic of Dcr-2 processing of primary hpRNA (pri-hpRNA) transcripts into 21-22 nt siRNAs. pri-hpRNAs can be detected using RNA-seq, and the corresponding siRNAs derived from the hairpin can be identified via small RNA-seq. (B) *hpRNA1* illustrates the behavior of hpRNA-derived testis RNA products in wildtype and *dcr-2* mutants. Normalized RNA-seq and spike-in normalized sRNA-seq tracks of the hpRNA shown in *w[XD1]* and *dcr-2*. The top two tracks show RNA-seq of primary hpRNA transcript is increased in *dcr-2* mutants due to upregulation of hpRNA primary transcript. The middle tracks show that the 5’ and 3’ ends of *pri-hpRNA1* are defined by 5’-seq and 3’-seq data, respectively. The bottom two tracks show that *hpRNA1*-derived small RNAs are biased to 21 nt and mostly eliminated in *dcr-2* mutant.

Together, this combination of datasets permits functional categorization of genuine hpRNAs with high specificity, as illustrated by *hpRNA1* (**Figure 1B**) and others (**Supplementary Figure 3**).

Strikingly, on the genomewide scale, hpRNA precursors were amongst the highest-upregulated transcripts in *dcr-2* mutants (**Figure 2**). We initially assessed this using the set of *Dmel* hpRNAs, all of which are conserved in *Dsim* (Wen *et al*., 2015). In MA plots of gene expression in *Dmel* and *Dsim dcr-2* testis compared to their respective genetic controls (**Figure 2A-D**), the known hpRNAs dominate the highest-upregulated transcripts in *dcr-2* mutants of both species. For example, *Dmel*-*hp-mir-997-1* was the 2nd-highest elevated locus genomewide, and 9/10 hpRNAs were in the top 35 upregulated loci in *Dmel* (**Figure 2A-B**). We similarly observed that pri-hpRNAs of *Dsim* orthologs of *Dmel* hpRNAs were highly elevated in *dcr-2* mutants (**Figure 2C-D**). These effects were specific, since primary miRNA transcripts were largely unaffected in *dcr-2* mutants of either species (**Figure 2B, D**). This was expected, as miRNAs are processed instead by Dcr-1. One exception was *mir-985*, whose primary transcript was elevated in mutants of *Dmel dcr-2* (**Figure 2A-B**), but not in *Dsim dcr-2* (**Figure 2C-D**). The reason for this discrepancy is unknown, but a potential explanation is that transcription of *mir-985* is elevated in *Dmel* as a secondary effect that is not shared in *Dsim*. Finally, hpRNA-derived siRNAs were strongly depleted in *dcr-2* mutants, including from all previously classified *Dmel* hpRNAs and their *Dsim* orthologs (**Figure 2E-F**).

**Figure 2.**
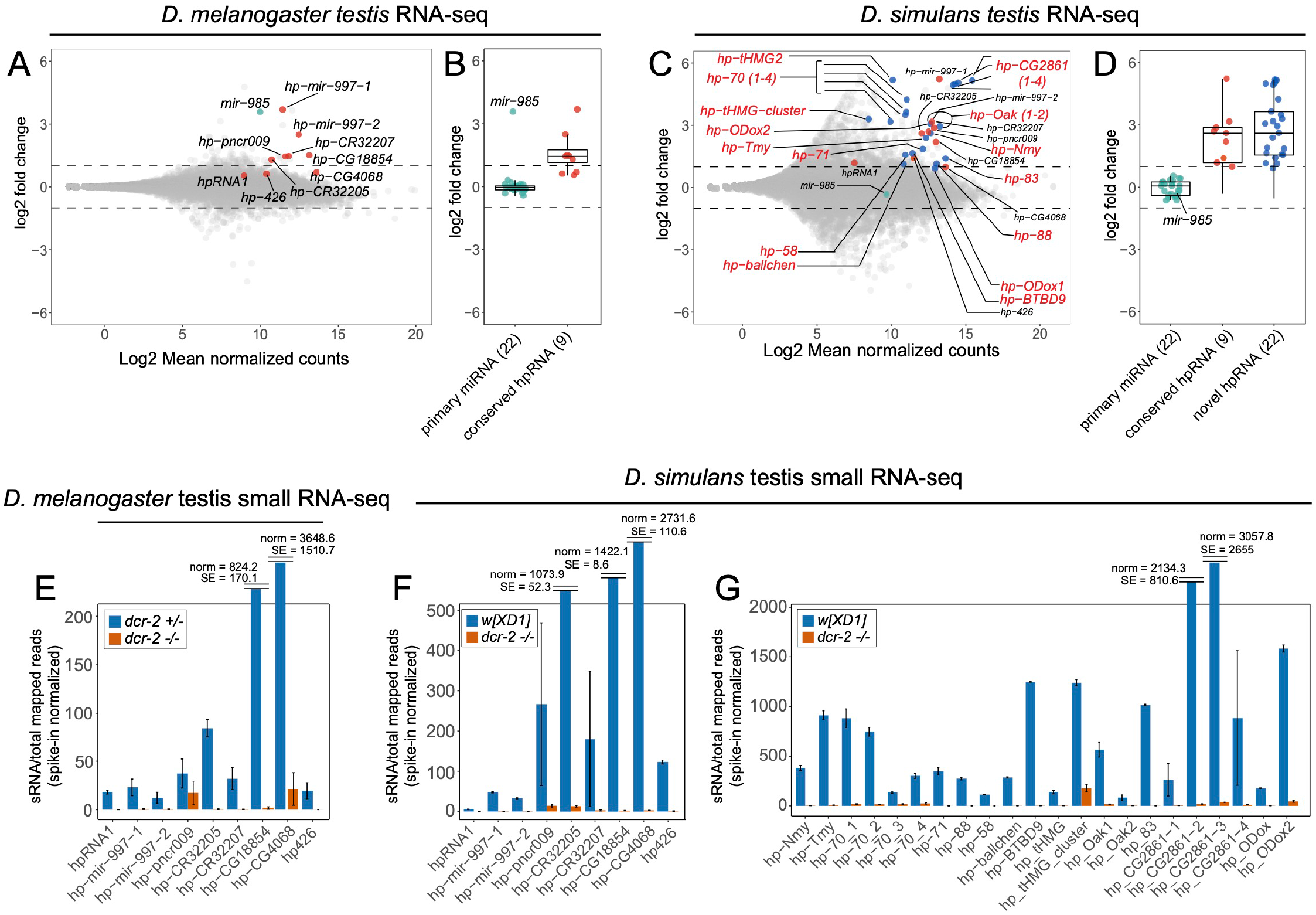
Comparative hpRNA annotation using functional genomic data. (A) MA-plot comparing *Dmel* wild-type and *dcr-*2 testis RNA-seq data. Orange dots denote hpRNAs conserved with *Dsim*, including one newly-recognized locus *hp-426*. (B) Primary hpRNA transcripts are all elevated in *Dmel dcr-*2 testis, unlike primary miRNA transcripts; *mir-985* is a lone exception. (C) MA-plot comparing *Dsim* control *w[XD1]* and *dcr-2* testis RNA-seq data. Orange dots mark hpRNAs conserved with *Dmel*, while blue dots indicate *de novo* hpRNAs in *Dsim*. (D). Comparison of primary hpRNA and primary miRNA transcripts in *Dsim*. All pri-miRNA transcripts are unchanged (including *mir-985*), while all pri-hpRNA transcripts are elevated in both conserved and de novo hpRNAs. (E-G) Expression of hpRNA-derived small RNAs in *Dmel* (E) and *Dsim* (F-G). All hpRNA-siRNAs are decreased in *dcr-2* mutants, in both species and regardless of the hpRNA age.

Inspection of the local genomic regions of known hpRNAs revealed provocative differences between *Dsim* and *Dmel* in the vicinity of hpRNA clusters, and provided additional evidence for rapid flux in hpRNA loci. The largest set of dispersed hpRNA loci in *Dmel* are members of the *hp-pncr009* family, for which 3 separate hpRNAs (*hp-pncr009*, *hp-CR32207*, and *hp-CR32205*) are interspersed with 9 protein-coding target genes of the *825-Oak* family (Wen *et al*., 2015). We have theorized that transcription across pairs of divergently-oriented *825-Oak* family loci might beget *pncr009* hpRNAs. Interestingly, in the short evolutionary distance that separates *Dmel* and *Dsim*, we identify two additional *pncr009-* class hpRNAs in *Dsim* (**Supplementary Figure 4A**). To facilitate intuitive connection of these hpRNAs to target genes in the *825-Oak* family, we named these novel *Dsim* hpRNAs as *hp-Oak1* and *hp-Oak2* (**Figure 2C-D**).

We also documented evolutionary flux in the tandem hpRNA repeats of the *hp-CG4068* cluster. Although the copy number was potentially in question from prior short-read genome assemblies, the recent availability of *simulans*-clade PacBio genomes (Chakraborty et al., 2021) demonstrates radical copy number of the *hp-CG4068* cluster. There are 20 tandem copies in *Dmel* but only 9 tandem copies in *Dsim* (**Supplementary Figure 4B**), as well as 14 copies in *Dmau*, and 10 copies in *Dsech* (not shown). Overall, there is high evolutionary divergence in the copy number of hpRNAs located in both genomically linked copies (*hp-pncr009* cluster) as well as in tandem copies produced from a common transcript (*hp-CG4068* cluster). Such dynamics are much greater than observed for canonical miRNAs, which only occasionally exhibit similar changes amongst these species (Mohammed et al., 2014a; Mohammed et al., 2014b).

### Unidirectional expansion of hpRNAs in *Dsim* compared to *Dmel*

Our small RNA and RNA-seq testis datasets robustly validated known hpRNAs, and yielded candidate evidence for distinct hpRNA content between *Dmel* and *Dsim*. Thus, we undertook more comprehensive hpRNA annotations as a basis of more comprehensive evolutionary comparisons. In particular, we sought highly structured loci that produce ~21 nt-biased, Dcr-2-dependent small RNAs, that also accumulate primary transcripts in *dcr-2* mutants (**Figure 1A**). However, we did not absolutely require pri-hpRNA changes in *dcr-2*, for several reasons. First, pri-hpRNA transcripts might not be stable and/or might be subject to other RNA decay pathways. Taking the canonical miRNA pathway as an analogy, not all pre-miRNAs accumulate in mutants of cytoplasmic Dicer, and not all pri-miRNAs accumulate as unprocessed transcripts in mutants of nuclear Drosha. Second, it was conceivable that some loci are processed earlier in development to yield stable small RNAs, but are not substantially transcribed in adult testis. Finally, transcription of some pri-hpRNA loci might be decreased in RNAi mutants.

In *Dmel*, we previously annotated hpRNAs from nearly 300 small RNA libraries of diverse developmental, tissue and cell origins, yielding only nine confident hpRNAs (Wen *et al*., 2015). Given that testis is the predominant location of hpRNA-siRNA accumulation, we interrogated our new genetically paired testis data for evidence of additional Dcr-2-dependent inverted repeat loci. However, beyond previously known hpRNAs, we only recovered a single new hpRNA, *hp426* (**Figure 2A-B** and **2E**), which produces siRNAs in *Dsim*. Thus, we did not identify any *Dmel*-specific hpRNAs.

A very different picture emerged from analysis of *Dsim*. Although we had far fewer small RNA libraries to annotate from, especially of testis datasets, we found numerous novel hpRNAs. These were highly confident since nearly all of them exhibited reciprocal behavior of primary transcripts and mature siRNAs, when comparing control and *dcr-2* testis libraries (**Figure 2C-D** and **G**). In fact, most novel pri-hpRNA loci resided amongst the most-upregulated transcripts across *Dsim dcr-2* testis (**Figure 2C**). A majority of these accumulated discrete spliced RNAs. However, depending on the locus, transcript coverage was often non-uniform. This was particularly the case within highly-duplexed portions of foldback structures (**Supplementary Figure 5**), evidently indicating that library construction was compromised within highly double-stranded transcript regions. Coverage was also an issue at transcript termini, which is generally the case with typical RNA-seq protocols (Shenker et al., 2015).

We therefore employed two more approaches to help annotate pri-hpRNA transcripts in control and *dcr-2* mutant testis: analysis of 5’ ends from low-input RNA (Schon et al., 2018) and 3’-end sequencing to determine polyadenylation sites (Sanfilippo et al., 2017). As illustrated in **Figure 1B**, 5’-seq and 3’-seq directly visualize the beginnings and ends of pri-hpRNA transcripts, and indicate that hpRNA progenitors are mRNAs. We note that some hpRNA loci are exact genomic copies that remain valid upon scrutiny of the highly contiguous PacBio *Dsim* assembly (Chakraborty *et al*., 2021), a phenomenon we return to later in this study. However, even when conservatively counting identical hpRNA copies within a tandem cluster as a single locus, there are 23 distinct, *de novo* hpRNA transcription units in *Dsim*, which can be assigned to 15 families that are not simply genomic copies (**Figure 2C-D, G**). As described later, some of these families can still be traced as sharing evolutionary heritage (as is the case for Nmy/Tmy, which are related in sequence but have separable functions).

Overall, the radical and asymmetric expansion of hpRNAs in *Dsim* compared to *Dmel* strongly suggests that the RNAi pathway has been deployed adaptively in these sister species, putatively to address emerging regulatory situations such as intragenomic conflicts.

### Target network of *Dsim*-specific hpRNAs

We next investigated targets of novel *Dsim* hpRNAs. In our previous work in *D. melanogaster*, we observed that hpRNAs typically exhibit substantial complementarity to one or a few target genes, ranging from an individual siRNA to extended regions that encompass multiple siRNAs (Okamura et al., 2008; Wen *et al*., 2015). On this basis, we proposed that hpRNAs typically derive from their targets (Wen *et al*., 2015), analogous to plant miRNAs (Allen et al., 2004). This set the stage that it seemed plausible, if not likely, that de novo hpRNAs in *Dsim* might also have overt complementary targets. Indeed, we were able to identify compelling targets with antisense matching to most newly-recognized *Dsim*-specific hpRNAs. In the following sections, we describe notable insights from specific aspects of the *Dsim* hpRNA target network.

### Unexpected complexity in the *Dsim* hpRNA network related to Dox family loci

We recently reported that multiple X-linked Dox family genes, including two newly-recognized members (*PDox1* and *PDox2*), share an HMG box domain that is derived from protamine (Muirhead and Presgraves, 2021; Vedanayagam *et al*., 2021). This was notable as the replacement of histones by protamines comprises a key transition in the normal condensation of paternal chromatin during sperm maturation (Rathke et al., 2014; Wang et al., 2019). The Dox family members are targeted by newly-emerged hpRNAs (e.g. Nmy and Tmy), and both Dox family genes and their complementary hpRNAs have undergone substantial expansion in *Dsim*-sister species (Lin *et al*., 2018; Muirhead and Presgraves, 2021; Vedanayagam *et al*., 2021). These data reflect an ongoing and intense intragenomic conflict within the *simulans*-clade. Now, with *Dsim* hpRNA target maps based on functional genomics, we reveal additional, *de novo* innovations within the Dox/hpRNA regulatory network.

We recently identified a sub-lineage of Dox-related loci that lack the HMG box (Vedanayagam *et al*., 2021). Via synteny comparisons with *D. melanogaster*, we inferred these to derive originally from fusion of a protamine-like copy in between *CG8664* and *forked* loci in an ancestor of *simulans* clade species, termed “original Dox” (*ODox*, **Figure 3A**). Our evolutionary tracing supports that *ODox* spawned the contemporary *Dox* family genes *PDox*, *UDox*, *MDox* and *Dox* across *simulans* clade Drosophilids (Vedanayagam *et al*., 2021). Perhaps confusingly, then, the *ODox* locus in contemporary *simulans*-clade species retains segments of *CG8664* and the 5’ UTR of *protamine*, but has lost its HMG box (**Figure 3B**). *ODox* subsequently duplicated and mobilized to yield the related *ODox2* locus, which shares predicted domains with *ODox* and lacks an HMG box, but also has divergent sequence material (**Figure 3B**). *ODox* and *ODox2* loci are proximal to centromere on the X chromosome at ~16Mb on the *Dsim* long-read assembly (Chakraborty *et al*., 2021), while the contemporary amplification of *Dox* family genes occurred at a distal genomic window of ~9-10Mb (**Figure 3B**).

**Figure 3.**
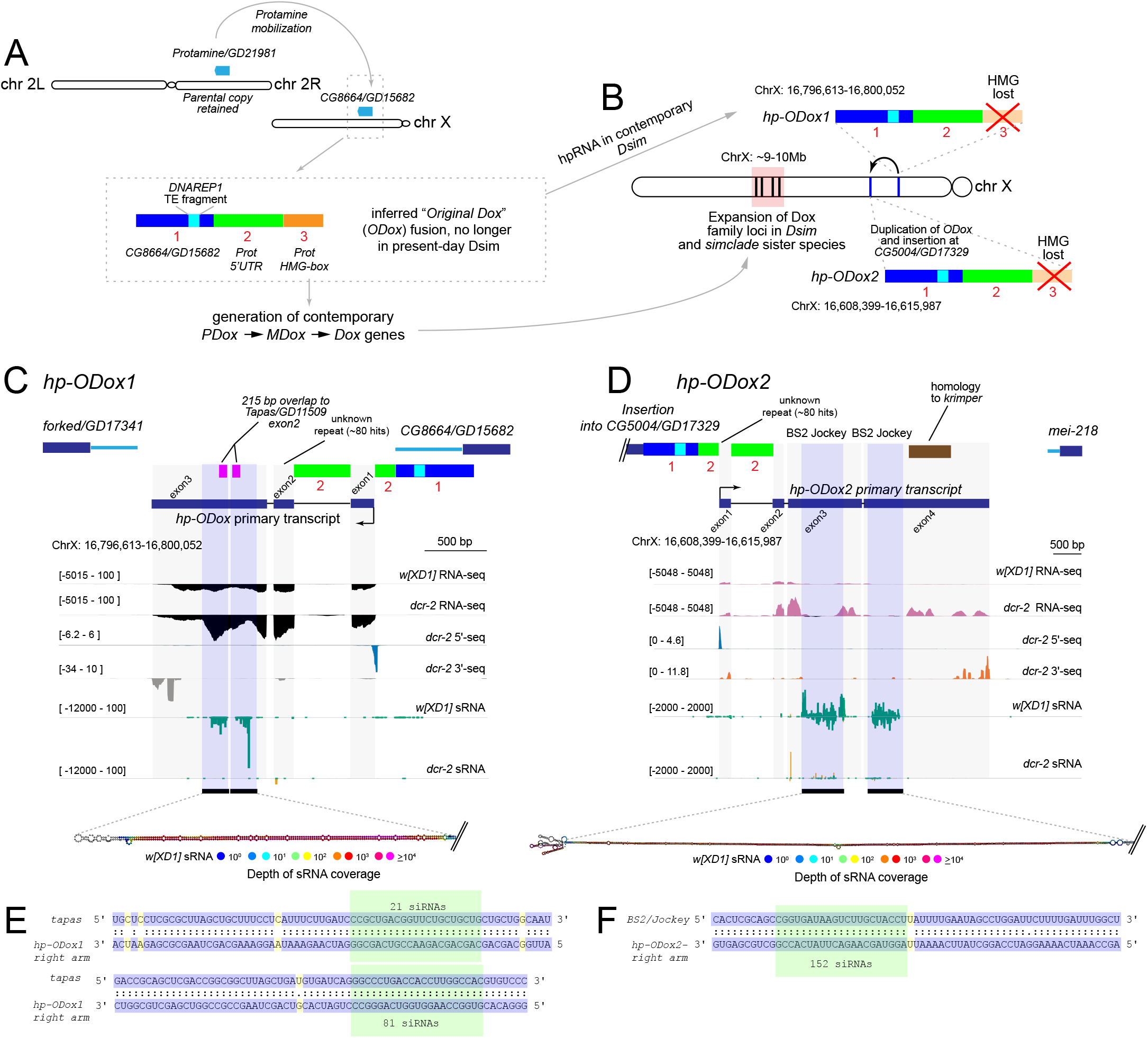
Innovations in the Dox-related hpRNA network. Schematic of the evolution of *Dox* family genes, and the contemporary state of inferred, ancestral fusion of *Protamine* and *CG8664* (*Prot-CG8446* fusion) that birthed the *Dox* family genes. Segments 1, 2 and 3 in the inset box (inferred *ODox* fusion), shows genomic regions corresponding to their sequence of origin. Segment 1 (blue) is derived from *CG8664*, segment 2 (green) is derived from the 5’ UTR of *Protamine*, and segment 3 (orange) contains *Protamine* coding sequence, including the HMG-box domain. Within segment 1, is a fragment of *DNAREP1* TE (turquoise). While contemporary Dox family genes *Dox*, *MDox*, and *PDox* share this segment structure, we infer that the ancestral “*ODox*” locus lost the HMG segment in contemporary *Dsim* (B). In contemporary *Dsim*, the ancestral *ODox* fusion is an hpRNA (*hp-ODox1*) and its duplication and insertion at *CG5004/GD1739* hosts another hpRNA (*hp-ODox2*). (C) *hp-ODox1* is a discrete hpRNA locus, revealed by upregulation of its primary hairpin transcript in *dcr-2* RNA-seq, presence of Dcr-2-dependent siRNAs, and demarcated by 5’-seq and 3’-seq. The inverted repeat arms of *hp-ODox1* bear homology to *Tapas* (pink), and other regions of *hp-ODox1* bear TE sequences. (D) *hp-ODox2* bears all the genomic signatures of an hpRNA, and its inverted repeat arms are homologous to *BS2/Jockey* TE. *pri-hp-ODox2* also bears homology to *Krimper*, although this sequence is not part of the inverted repeat and does not yield siRNAs. (E) Sequence alignment between *hp-ODox* and *tapas*. Examples of siRNAs with fully complementarity to *tapas* are boxed in green. (F) Antisense complementarity between *BS2/Jockey TE* and *hp-ODox2* with examplar siRNAs highlighted in green.

Given that HMG box domains are likely central to distorting function of Dox family factors (Muirhead and Presgraves, 2021; Vedanayagam *et al*., 2021), we wondered about the functional relevance of Dox-superfamily loci lacking HMG Box domains.

Unexpectedly, we realized that both *Dsim ODox* and *ODox2* contain inverted repeats that bear functional hpRNA signatures, i.e., they generate Dcr-2-dependent small RNAs and their primary transcripts are upregulated in *dcr-2* mutants (**Figure 2, Figure 3C-D**). Accordingly, we renamed these loci *hp-ODox1* and *hp-ODox2*. Detailed analysis reveals further unanticipated features of their domain content. In particular, the inverted repeat at *ODox* bears ~200 bp homology to exon 2 of *Tapas*/*GD11509* and generates siRNAs with antisense complementarity to the parental *Tapas*/*GD11509* on chr2R (**Figure 3E**). In addition, *hp-ODox1* transcript contains a mix of repetitive sequences (**Figure 3C**).

Unexpectedly, the hairpin at *hp-ODox2* is not homologous to the hairpin in *hp-ODox1*, despite their shared lineage. Instead, the *hp-ODox2* inverted repeat contains sequence from the *BS2/Jockey* transposable element (TE), generating siRNAs with antisense complementarity to *BS2/Jockey* (**Figure 3F**). In addition, other regions of the *pri-hp-ODox2* transcript bear other repetitive sequences as well as fragments from *Krimper*. However, the inverted repeat does not include *Krimper* sequence, and *Krimper*-targeting siRNAs were not observed.

These observations are intriguing, since Tapas and Krimper are both piwi-interacting RNA (piRNA) factors involved in transposon suppression in the female germline (Lim and Kai, 2007; Patil et al., 2014), and both *hp-ODox1* and *hp-ODox2* have also incorporated numerous other TE fragments. We note the Presgraves group identified homology between these loci (termed X:17.1 and X:17.2) and *Tapas*/*Krimper*, respectively (Muirhead and Presgraves, 2021), but these loci were not recognized as inverted repeat containing hpRNAs. Our analyses indicate that derivatives of an ancestral “*ODox*” gene now generate hpRNA-siRNAs in contemporary *Dsim*, and are presumably engaged in distinct genetic conflicts (**Figure 3**). Overall, the genetic conflict initially augured by the identification of *Dox* and *Nmy* (Tao *et al*., 2007a; Tao *et al*., 2007b) is actually embroiled in a far more extensive and rapidly evolving network of putative meiotic drivers and suppressors, and may in fact integrate activities of the siRNA and piRNA pathways. Relevant to this, the third recognized *sex ratio* meiotic drive system in *Dsim* (“Paris”), is driven by HP1D2 (Helleu *et al*., 2016), a derivative of the core piRNA factor Rhino (Klattenhoff et al., 2009; Mohn et al., 2014). Thus, there are recurrent linkages of piRNA factors to *sex ratio* meiotic drive, notwithstanding that TEs are themselves selfish genetic elements.

### Innovation of *Dsim* hpRNAs that target other HMG-Box loci

A related branch of *Dsim* hpRNAs and targets involves multiple tHMG loci. *Drosophila* testis specifically expresses numerous HMG box factors, some of which fall into the “MST-HMG box” subclass (Doyen et al., 2013). *Drosophila* protamines belong to this class, and given that HMG-Box domains of Dox family loci are most closely related to protamine (Muirhead and Presgraves, 2021; Vedanayagam *et al*., 2021), they can also be classified as MST-HMG box members. tHMG is not included in the MST-HMG box family, but is nevertheless also testis-specific and expressed at highest levels during the histone-to-protamine transition (Gartner et al., 2015). We note evidence for rapid evolution of testis HMG box loci, since protamine is locally duplicated in *Dmel* (*MST35Ba/Bb*), but exists as a single copy in simulans clade species (Doyen *et al*., 2013; Vedanayagam *et al*., 2021). Similarly, tHMG is locally duplicated in *Dmel* (*tHMG1* and *tHMG2*), but bears a single copy in the syntenic region of *Dsim* (**Figure 4A**).

**Figure 4.**
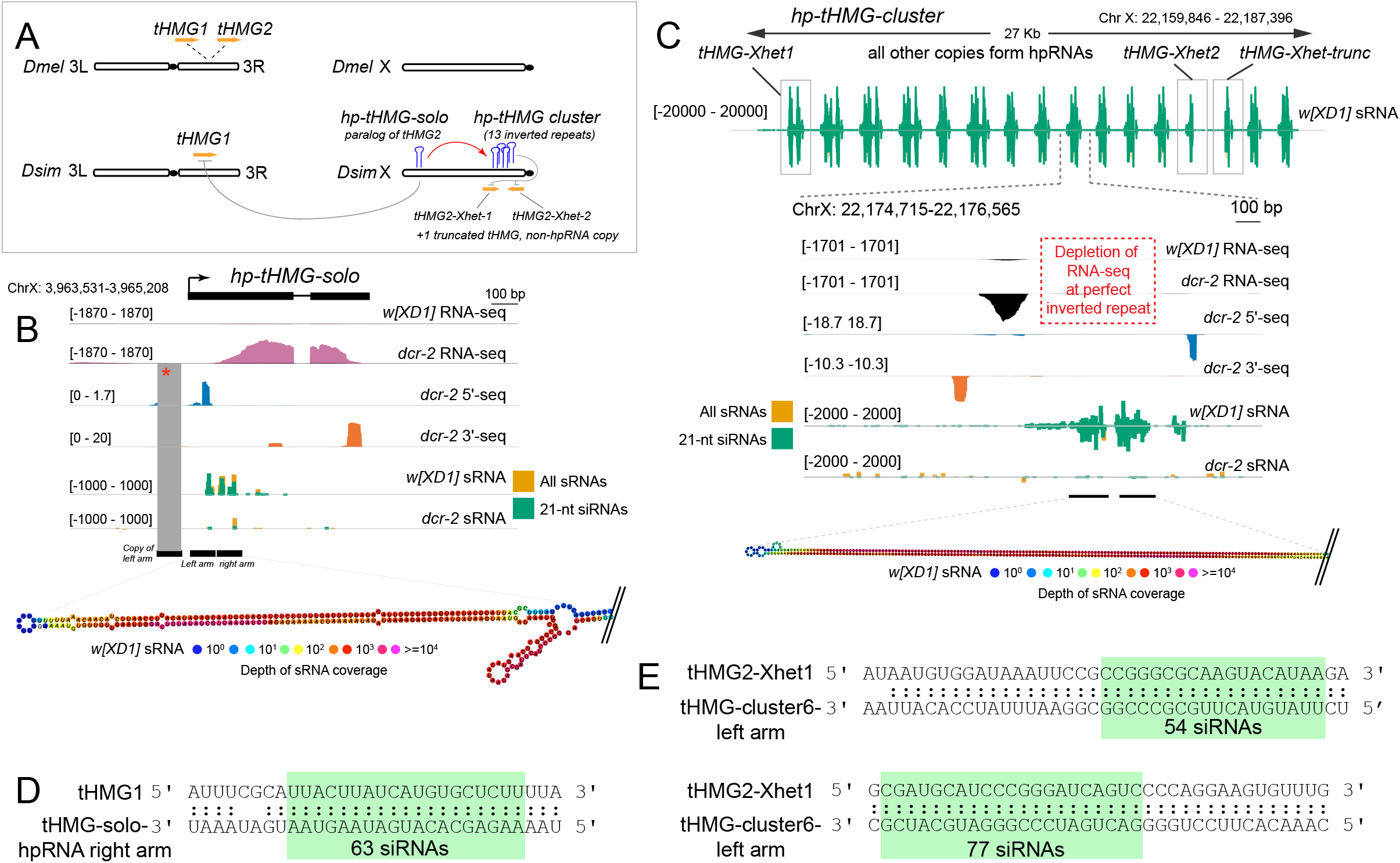
Novel *Dsim* hpRNAs related to testis HMG Box (tHMG) loci. (A) Schematic of tHMG loci in *Dmel* and *Dsim*, and the origins of hpRNA suppressors and targets encompassing multiple novel X-linked tHMG-derived loci. tHMG is locally duplicated in *Dmel*, but the syntenic location of *Dsim* bears a single gene. However, *Dsim* X chromosome bears multiple *de novo* hpRNAs with homology to tHMG, a single copy locus and another genomic cluster containing 13 tandem hpRNAs. (B) Genomic details of the *hp-tHMG-solo* locus, which is a spliced transcript with defined 5’ and 3’ ends, which generates abundant small RNAs from its duplex arm. Note that the 5’ region of *hp-tHMG-solo* has been locally duplicated. (C) Genomic details of the *hp-tHMG-cluster* locus. This region bears 13 nearly identical hpRNAs, although the nature of the primary transcript(s) is not clearly evident, although 6 cluster units show evidence for 5’-seq and 3’-seq (detailed in **Supplementary Figure 6**). RNA-seq data across the tHMG-cluster locus appears depleted in the hpRNA duplex regions. Small RNA tracks show Dcr-2-dependent siRNAs from each cluster member. (D, E) Examples of antisense complementarity between tHMG genes and hp-tHMG-siRNAs, including autosomal *tHMG1* and *hp-tHMG-solo*, and *tHMG-Xhet1* and tHMG cluster loci.

Our functional profiling of *Dsim* RNAi mutant testis allows us to discern further evolutionary dynamics of HMG box-related hpRNAs (**Figure 4A**). First, we find that *Dsim* contains a locus related to autosomal *Dmel tHMG2*, which has mobilized to the X chromosome (**Figure 4A**). Detailed inspection shows that this locus is actually an hpRNA, as it generates Dcr-2-dependent small RNAs, and is associated with a spliced transcript that is derepressed in *dcr-2* mutants (**Figure 4B**). As is the case with certain other strong inverted repeat loci, the RNA-seq signal is poorly represented in the duplex arms of the hairpin, which resides near the 5’ end of the primary transcript. However, the presence of capped and polyadenylated species provide experimental evidence for its termini. Of note, there is a local tandem duplication of the left arm of the hairpin, including within 5’-end mapping data (**Figure 4B**), and hints that its genesis as an hpRNA involved local duplications of a transposed sequence.

There are further complexities. We observe recent amplification of *tHMG2* in the heterochromatin boundary of chromosome X (Xhet), associated with numerous tandem hpRNAs within ~27kb (*hp-tHMG-cluster*) (**Figure 4C**). Based on the local duplication of *tHMG2*, and hpRNA secondary structure, we classify the 13 *hp-tHMG-cluster* hpRNAs into 3 subcategories (**Supplementary Figure 6**). The *hp-tHMG-cluster* also harbors three paralogs of *tHMG2* that are not part of inverted repeats. Of these, two paralogs appear to be full-length copies (88aa, compared to *Dsim-tHMG2* ortholog at 91aa), while a truncated paralog within this cluster encodes for only 24 aa (**Figure 4C and Supplementary Figure 6**).

At present, the nature of the primary transcript for the *hp-tHMG* cluster is uncertain, as near perfect complementarity of inverted repeats results in depletion of RNA-seq signal within *tHMG* hpRNA copies. We document 13 local inverted repeats (hpRNA copies) within the *hp-tHMG-* cluster. Of these, 6 have flanking 5’-end and 3’-seq signals, along with RNA-seq mapping between the hairpin arms that is upregulated in *dcr-2* mutants. While it is unclear how many individual transcription units exist with this hpRNA cluster, it is clear that the solo and clustered tHMG hairpin loci (located on opposite ends of the *Dsim* X chromosome) are evolutionarily related, to each other and to the parental autosomal copy of tHMG (**Figure 4C** and **Supplementary Figure 6**). Indeed, we identify abundant siRNAs generated from *hp-tHMG-solo* and the *hp-tHMG-cluster* that exhibit perfect complementarily to autosomal *tHMG1* and/or novel paralogs of *tHMG2* on Xhet (**Figure 4D**).

From these data, several themes emerge in the ongoing co-evolutionary arms race between hpRNAs and their HMG-box targets. First, at least two sub-families of HMG-box loci in testis (protamine and tHMG) exhibit rapid dynamics in copy number and location, even between very closely-related species. Second, members of both sub-families are suppressed by RNAi, revealing recurrent association of HMG-box loci with intragenomic conflict. Third, both examples include amplification of X-linked HMG-box copies. Together, these data suggest that repeated amplification of HMG-box loci are genetic weapons in a sex chromosome conflict, which triggers the emergence of concomitant hpRNA suppressors to silence meiotic drive.

### Delineation of stages of hpRNA emergence and evolution

With broader evidence that hpRNAs typically target specific genes, or groups of related loci, we sought broader perspective on evolution of hpRNA regulatory networks. Although all *Dmel* hpRNAs emerged relatively recently (Wen *et al*., 2015), it is instructive to note that all *Dmel* hpRNAs are conserved in *Dsim* (**Figure 2E-F**). Therefore, we may consider *Dmel* hpRNAs to be relatively old, compared to the numerous *de novo Dsim* hpRNAs we identify in this study (**Figure 2C,G**). In principle, then, this collection of extremely young *Dsim* hpRNAs may illuminate the earliest stages of hpRNA birth.

An ongoing conundrum concerns “where” hpRNAs come from, especially as the functionally validated hpRNA-target relationships comprise examples where the hpRNA is genomically distant from its target (Czech et al., 2008; Lin *et al*., 2018; Okamura *et al*., 2008; Wen *et al*., 2015). This could be due to mobilization of the hpRNA locus itself, or to derivation of the hpRNA via retrogene insertion (Tao *et al*., 2007a; Tao *et al*., 2007b). In the latter case, acquisition of a promoter may be an issue. We note that formation of *Dsim hp88* occurred with the 3’ region of an existing apparent non-coding locus *CR43306* (**Figure 5A**), suggesting that hpRNAs could take advantage of pre-existing transcription units for their expression.

**Figure 5.**
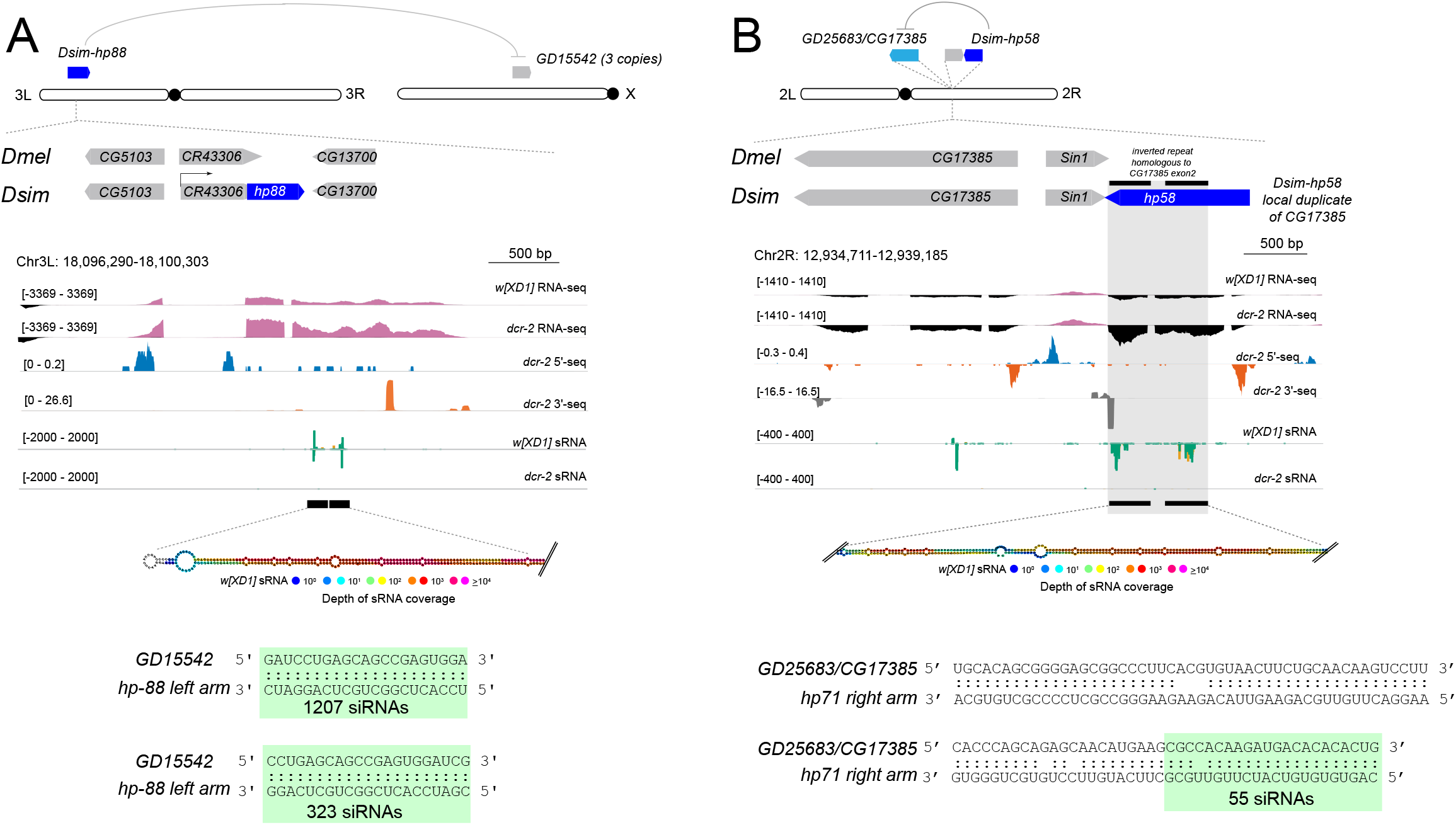
Genomic features of novel *Dsim* hpRNAs inform early stages in their birth. (A) *Dsim*-hp88 emerged within the 3’ UTR of non-coding RNA CR43306, which is syntenic between *Dmel* and *Dsim*. The inverted repeat arms bear homology to *de novo* paralogs on the X chromosome *GD15542* (3 copies expressed, but there 13 other paralogous sequences of this novel gene on the X). *GD15542* is upregulated in *dcr-2* mutant indicating functional suppression via RNAi. hpRNA-target alignments for both *Dsim-hp58* and *Dsim-hp88* are provided with highlighted green box showing fully complementary siRNAs to their targets, *CG17385* and *GD15542*, respectively. (B) *Dsim-hp58* is a *de novo* hairpin that was born one gene away from its progenitor gene *CG17385*. It bears an inverted repeat fragment of exon 2 of *CG17385*. Similar to *pri-hp58*, *CG17385* is upregulated in *dcr-2* mutant testis RNA-seq data, indicating its functional suppression via the RNAi pathway.

Our catalog of *de novo* hpRNAs includes many other hpRNA loci that are genomically distant from their targets. However, the precedent of the interweaved locations of multiple related hpRNAs and targets of the *hp-pncr009/825-Oak* families in *Dmel* suggested that some hpRNAs might be born from a genomic location close to their target. We now find several examples of this. For example, *Dsim hp58* is a newly-emerged hpRNA that is adjacent to its pre-existing target gene *GD25683/CG17358* (**Figure 5B**). Similarly, we identify a novel hpRNA located at *Dsim ballchen* (*GD18116*), created by a partial duplication of the 5’ region adjacent to the conserved *ballchen* gene annotation. This is reminiscent of the partial 5’ duplication of the *hp-tHMG-solo* locus (**Figure 4**).

We also emphasize that hpRNA evolution frequently involves emergence of multiple copies. Beyond the described examples of expanding *hp-pncr009* family clustered loci and *hp-CG4068* tandem hpRNAs, we discovered novel amplified hpRNAs in *Dsim*. These include tandem duplicates (as in the novel tHMG-related hpRNA cluster), local genomic duplicates of independent transcription units (as in de novo *hp-pncr009* family members, **Supplementary Figure 4B**), or genomically dispersed copies (as in the case of four copies of the *hp70* family, **Figure 5B**). Finally, we highlight *Dsim hp88*, which was appended to an existing non-coding locus, but contains genetic material from the trio of *GD15542* gene copies located on another chromosome (**Figure 5B**), much like amplifications of Dox family genes trigger hpRNA birth (Muirhead and Presgraves, 2021; Vedanayagam *et al*., 2021). The frequency of amplified hpRNAs and targets seems analogous to the notion of “molecular accordions” during viral arms races, where transient expansion of a selfish gene product was proposed to facilitate its exploration of sequence diversity during escape from host defense (Elde et al., 2012). Here, the existence of amplified hpRNA loci of a given family may also aid the process of co-evolution with target genes.

### Functional repression by hpRNAs is most overt for the youngest siRNA loci

With an expanded view of hpRNAs and targets in hand, we addressed the larger consequences of RNAi loss on the testis transcriptomes of both species. Interestingly, although cytological consequences of RNAi loss on spermatogenesis is substantial in *Dmel* (Wen *et al*., 2015) and severe in *Dsim* (Lin *et al*., 2018), their respective transcriptome responses were relatively restricted (**Figure 2A, C**). For example, in *Dmel*, only 108/16773 annotations detected >10RPM were 2-fold upregulated in *dcr-2* mutants compared to *dcr-2/+* heterozygous controls (FDR <1%). However, all hpRNAs (except *hpRNA1*, upregulated only 1.5-fold) were amongst the top 20 upregulated transcripts in *dcr-2* mutants, confirming their efficient metabolism by Dcr-2. hpRNA targets also responded directionally to RNAi loss, in that none were downregulated (**Figure 6**); however, many hpRNA targets were not detected in adult testis. Of the 8 conserved hpRNA loci between *Dmel* and *Dsim*, the *pncr009* cluster hpRNAs (*hp-pncr009*, *hp-CR32205*, and *hp-CR32207*) bear homology to 10 target genes belonging to the *825-Oak* family (Wen *et al*., 2015) (**Supplementary Figure 4**). In *Dmel dcr-2* mutant testis, all *825-Oak* family genes were <10RPM threshold, or indeed undetected. The *Dmel* orthologs of *825-Oak* loci are restricted to pupal gonads (http://flybase.org/), even though their corresponding hpRNA-siRNAs are detectable in adult testis. Of the remaining *Dmel* hpRNAs, *hp-mir-997-1* and *hp-mir-997-2* target *CG15040*, which registered only marginal directional change upon *dcr-2* loss. Only two genes, *ATP-synß* and *mus308*, targets of *hpRNA1* and *hp-CG4068* respectively, were elevated in *dcr-2* mutants (**Figure 6A**). The targets of *Dmel-Dsim* conserved hpRNAs exhibited similar expression profiles in *Dsim dcr-2* mutants (**Figure 6B**). Thus, there is functional repression of targets of the older, conserved hpRNAs in both species, but the effects are generally modest. Nevertheless, the regulatory effects on siRNA targets are still greater than with most miRNA targets (Agarwal et al., 2018).

**Figure 6.**
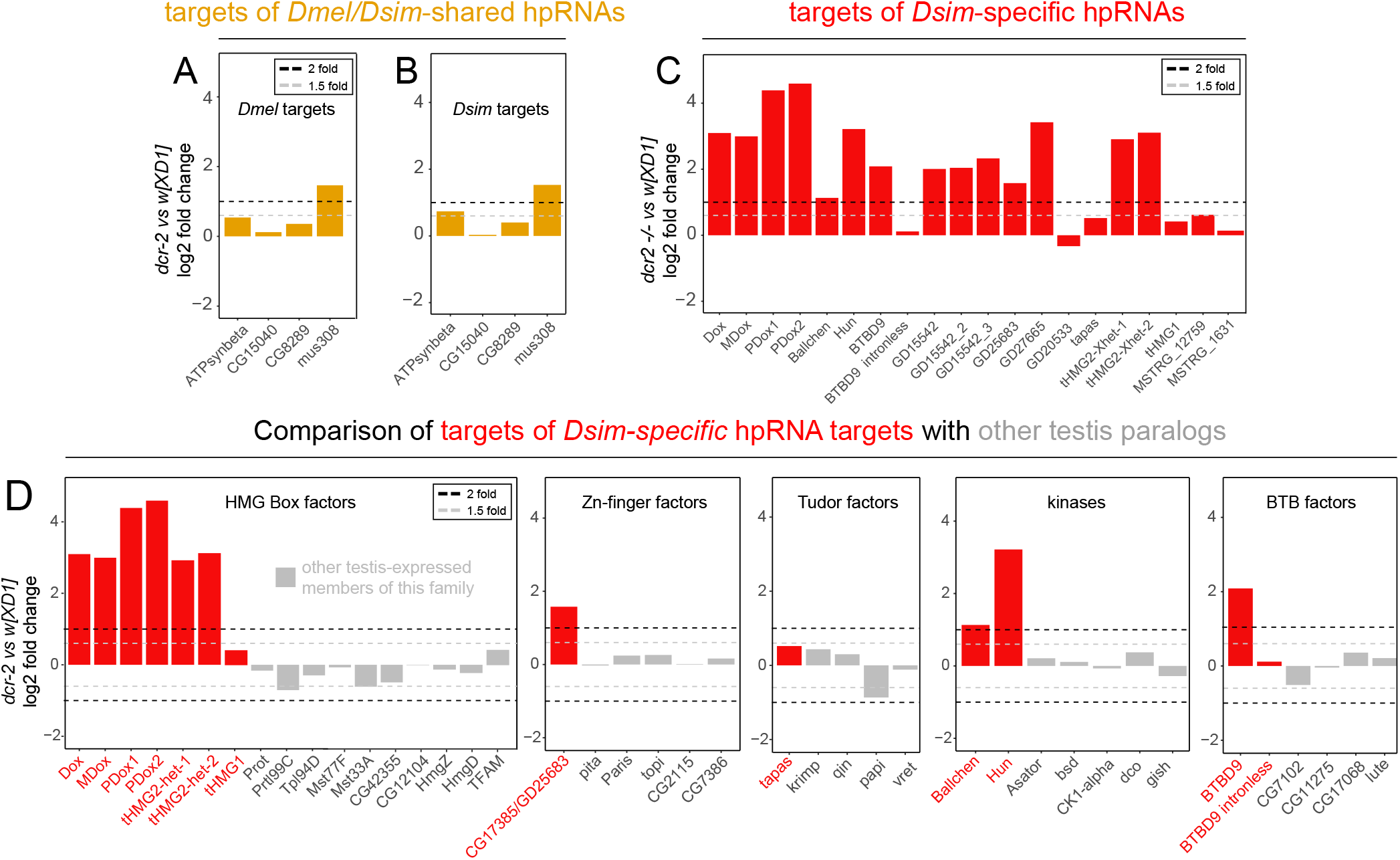
Preferential suppression of targets of young hpRNAs vs. older hpRNAs. Differential expression analysis of the targets of conserved and novel hpRNAs in *Dmel* and *Dsim* testis RNA-seq data. (A, B) Targets of hpRNAs conserved between *Dmel* and *Dsim*. Fold change values comparing wildtype and *dcr-2* mutant shown in yellow. Black and grey dotted lines show 2-fold and 1.5 fold changes in *dcr-2* mutant compared to wild type. Fold change values were estimated using two replicates each for the mutant and wildtype samples using DEseq package in R with an FDR < 1%. (C) Targets of *Dsim*-specific hpRNAs (red) and their fold changes in *dcr-2* mutant compared to wild type. Note the functional repression of targets of young hpRNAs is much greater than for older hpRNAs. For comparison, the expression changes of additional testis-expressed genes with similar domains as the hpRNA targets are shown in grey.

In striking contrast, the targets of the youngest siRNA loci, i.e. of *Dsim*-specific hpRNAs, showed substantial greater directional change upon *dcr-2* loss (**Figure 6C**). In *Dsim*, 976/15119 loci expressed >10RPM exhibit at least two-fold upregulation in *dcr-2* mutant compared to *w[XD1]* (FDR <1%). Of these, 22 novel and 7 conserved hpRNAs were in the top 200 derepressed genes. Moreover, 14/20 targets of novel hpRNAs in *Dsim* were among top 200 upregulated genes. To further assess whether the upregulation of *de novo* hpRNA targets is specific, we compared *de novo* hpRNA targets to related paralogs in *Dsim* that are also expressed in testis. For example, we detected directional change in expression of Dox family members upon loss of Dcr-2, but not their related HMG-box paralogs that are also expressed in testis (**Figure 6D**). Similar specificity was observed for other families of *de novo* targets (zinc-finger factors, Tudor proteins, kinases, and BTBD factors, **Figure 6D**).

Overall, the analysis of *Dsim* was particularly informative, since we could compare the properties of “young” and “older” hpRNA-target interactions. This was perhaps unexpected, since we had earlier used the latter cohort to derive clear evidence of adaptive co-evolution between hpRNAs and their targets (Wen *et al*., 2015). Due to insufficient orthologs, we lack statistical foundation to assess co-evolution with *simulans*-specific hpRNAs. Nevertheless, the picture is clear that younger hpRNAs mediate quantitatively greater target suppression than older hpRNAs. In particular, these data flip the rationale for small RNA mediated regulation relative to the miRNA pathway, for which recently-evolved loci might generally be neutral and only the oldest miRNAs appear to have biologically significant effects (Bartel and Chen, 2004).

### Biased X-linkage of *de novo Dsim* hpRNA targets, along with some hpRNAs, reflects likely roles in sex chromosome conflict

The “oldest” *Dsim* hpRNAs (i.e., still relatively young, but shared with *Dmel*), collectively target both young and ancient genes, which are distributed across all the chromosomes (with multiple targets on each of the larger chromosomes). Thus, there is no overt bias to the age and location of “old” hpRNA targets, beyond the fact that many hpRNAs and targets in the *hp-pncr009/825-Oak* network are clustered within a small genomic interval, and continue to expand actively (**Figure 7** and **Supplementary Figures 4, 8**).

**Figure 7.**
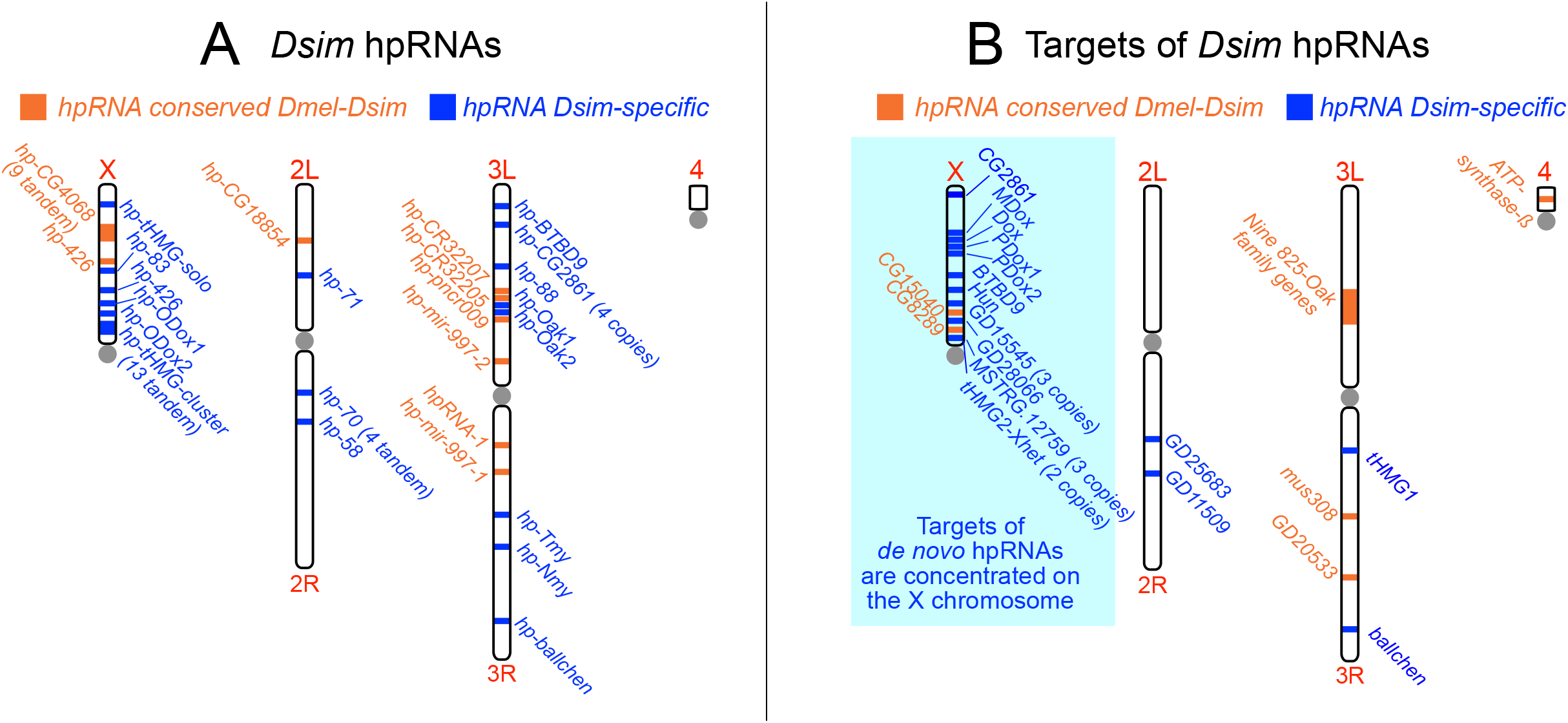
Biased location of *de novo* hpRNA targets on the *D. simulans* X chromosome. (A) Genomic location of *Dsim* hpRNAs, separated into ones that are conserved in *Dmel* (in orange) and ones that are *Dsim*-specific (in blue). Both classes of hpRNAs are distributed across all the major chromosome arms. (B) Genomic location of *Dsim* hpRNA targets, separated into ones whose hpRNA is conserved in *Dmel* (in orange) and ones whose cognate hpRNA is *Dsim*-specific (in blue). Note that the concentration of targets of *Dsim*-specific hpRNAs on the X suggests that they may comprise novel selfish genes involved in *sex ratio* meiotic drive.

On the other hand, a distinct pattern emerges with our collection of “young” *Dsim* hpRNAs. These very young hpRNAs (blue loci, **Figure 7A**) are distributed across the major chromosomes, in a pattern relatively similar to the older hpRNAs (orange loci, **Figure 7A**). On the other hand, the targets of these recently-emerged *Dsim* hpRNAs, exhibit a striking bias for X-chromosome localization (blue loci, **Figure 7B**). In addition to Dox and MDox, 14/20 (70%) other targets of *Dsim*-emerged hpRNAs are found on the X, whereas only 2/11 (18%) targets of hpRNAs shared with *Dmel* are found on the X. These genes are all testis-specific paralogs of older gene families, and many encode proteins with provocative roles that putatively relate to meiosis or chromosome segregation. As documented, these include four paralogous loci Dox, MDox, PDox1 and PDox2, which define a rapidly evolving set of young protamine-like gene copies with known or inferred meiotic drive activities (Lin *et al*., 2018; Muirhead and Presgraves, 2021; Tao *et al*., 2007a; Tao *et al*., 2007b; Vedanayagam *et al*., 2021). Moreover, we described in detail about the existence of additional X-linked tHMG loci, for which several protein-coding copies within the tHMG-cluster and the Xhet region are targeted by hpRNAs (**Figure 4** and **Supplementary Figure 6**).

*Hun Hunaphu* (*Hun*) is another intriguing X-linked hpRNA target. *Hun* is a newly-emerged X-linked derivative of *ballchen*, encoding a histone kinase that is essential for germline stem cell renewal (in both sexes) and meiotic chromosomal architecture (Cullen et al., 2005; Herzig et al., 2014; Ivanovska et al., 2005; Lancaster et al., 2007). However, like Dox family genes (Muirhead and Presgraves, 2021; Vedanayagam *et al*., 2021), *Hun* is a chimeric, young gene that has lost parts of Ballchen and gained new coding sequence (Arguello et al., 2006). Of note, Hun orthologs exhibit a large excess of nonsynonymous substitutions compared to Ballchen (Arguello *et al*., 2006). Ballchen is moderately upregulated in *dcr-2* mutant testis but *Hun* is much more strongly derepressed (**Figure 6**). In light of their strong suppression by endo-RNAi, we take this as a strong suggestion for a selfish function of Hun that necessitates silencing by a hpRNA.

Overall, the biased location of targets of *de novo* hpRNAs on the *Dsim* X broadens the scenario that the *Dsim* X is in genetic conflict with the Y. More importantly, our studies provide a molecular roadmap to directly identify putative selfish loci by virtue of their abundant siRNA-mediated repression. This unlocks the potential to identify the genes involved in otherwise cryptic intragenomic conflicts, for which cycles of drive and repression are poised to underlie speciation (Agren and Clark, 2018; Lindholm et al., 2016; Meiklejohn and Tao, 2010).

## Discussion

### Biological utilities of endogenous RNAi

It is now decades since the initial reports of RNA-based homology-dependent silencing (i.e., RNAi) (Fire et al., 1998; Napoli et al., 1990; van der Krol et al., 1990). The diverse experimental utilities of this pathway are well-established and continue to grow, and have finally given rise to several bona fide siRNA drugs (Zhang et al., 2021).

Nevertheless, the biological utilities of endogenous RNAi remain less-defined, especially for processes with phenotypic impacts (as opposed to simply perturbing small RNA biogenesis or gene expression). This is particularly the case for animal species that bear an independent, evolutionarily derived, RNAi pathway that is distinct from the ancestral miRNA pathway shared by all metazoans (Okamura and Lai, 2008).

For example, the endo-RNAi pathway mediated by core factors RDE-1 (Argonaute) and RDE-4 (dsRBD partner of Dicer) were amongst the first gene products found to be required for silencing by exogenous dsRNA triggers in *C. elegans* (Tabara et al., 2002). Their mechanistic involvement in an expansive regulatory hierarchy that generates a bewildering cascade of primary, secondary and even tertiary siRNAs, continues to be studied (Pak et al., 2012; Tsai et al., 2015). Nevertheless, the biological consequences of deleting these RNAi-specific factors (i.e., RDE-1 and RDE-4) have been largely elusive, beyond their failure to control viral infections (Lu et al., 2005). Notably, some recent studies provide evidence for the requirement of core endo-RNAi factors in transgenic models of pathogenic proteins (Long et al., 2014) or in the transgenerational inheritance of acquired traits (Alcazar et al., 2008; Devanapally et al., 2015; Kaletsky et al., 2020; Moore et al., 2019; Posner et al., 2019). These results extend themes in which nematode endo-RNAi is involved in suppressing aberrant and/or foreign transcripts, or in conveying epigenetic information analogous to, or even upstream, of the piRNA pathway.

In *Drosophila*, components of the dedicated RNAi pathway (i.e., Dcr-2 and AGO2) are similarly required for antiviral defense (Mondotte et al., 2018; Wang et al., 2006), and ongoing research has focused on mechanisms of siRNA biogenesis (Goh and Okamura, 2019; Lee et al., 2018; Sinha et al., 2018; Tsuboyama et al., 2018) and/or new regulatory arenas such as transcriptional silencing and dosage compensation (Deshpande and Meller, 2018; Lee *et al*., 2018; Nazer et al., 2018).

In this study, we expand on our recent proposal that a major utility of endo-RNAi in flies is to control intragenomic conflicts (Lin *et al*., 2018; Vedanayagam *et al*., 2021; Wen *et al*., 2015). This is conceptually similar to a role for endo-RNAi in suppressing transposable elements (Chung et al., 2008; Czech *et al*., 2008; Ghildiyal et al., 2008; Kawamura et al., 2008). Although the piRNA pathway is more well-known for TE suppression, the role of siRNA pathway in this process becomes more overt in certain sensitized conditions such as aging or neurodegeneration (Krug et al., 2017; Li et al., 2013), or when the piRNA pathway is compromised (Barckmann et al., 2018; van den Beek et al., 2018). However, TEs have certain gene expression signatures that help to signal their distinction from conventional genes (in particular, sense and antisense transcripts that are involved in biogenesis and targeting by siRNAs and piRNAs. On the other hand, hpRNA targets are distinct in that they are conventional mRNAs encoded by cellular genes, and they are not intrinsically involved in biogenesis.

### The hpRNA targeting landscape is diametrically opposite to miRNA regulation

We conceive two general classes of hpRNA targets. While hpRNAs are all evolutionary young, some of their targets are relatively old genes; e.g. targeting of *ATP synthase-ß* by hpRNA-1. We imagine there are adaptive reasons for why mild suppression of such loci imparts beneficial regulatory consequences. This may be due directly to the acquisition of elevated transcriptional properties of the targets, and/or by emergence of duplicated loci with preferred testis expression, which appears to be a relatively common process (Assis and Bachtrog, 2013; Kondo et al., 2017). In any case, the role for endo-RNAi here is to modulate target activity, since full suppression of these genes is clearly deleterious. Still, we may speculate further that as several well-conserved targets of hpRNAs encode protein activities that have been linked to speciation, such as heterochromatin, DNA damage, and energy homeostasis (Wen *et al*., 2015), these conserved genes may harbor selfish activities in certain species that warrants adaptive suppression by RNAi. Since the miRNA pathway does not typically does adaptive targeting, and instead relies upon capture of targets bearing invariant miRNA seed matches, the RNAi pathway may be more flexible to suppress such genes.

### Recurrent themes in RNAi-regulated networks: HMG Box factors, piRNA factors and TEs

Our annotation of hpRNAs in two species indicates that, as a rule, these endogenous siRNA loci comprise very short-lived genes. Thus, they can at best only mediate modestly conserved regulation. If this is the case, can we learn any general principles from such fast-evolving regulatory networks? In fact, when considering this study alongside recent literature, we find several recurrent themes that provide a framework for understanding how endogenous RNAi is harnessed in biology.

We recently found that *de novo* hpRNAs in the *simulans* clade are required to silence a newly-emerged, amplifying, and selfish set of X-linked protamine derivatives, namely the Dox family (Muirhead and Presgraves, 2021; Vedanayagam *et al*., 2021). Protamines are central factors that condense the sperm genome, and therefore seem ripe for co-option by selfish factors to disrupt paternal inheritance. Indeed, following removal of histones, multiple sperm nuclear basic proteins (SNBPs) play roles in packaging sperm chromatin, and most of these contain HMG box domains. We find that beyond the X-linked Dox family, there is a separate amplification of X-linked tHMG box genes in Dsim that are concomitantly associated with silencing by cognate hpRNAs. Thus, we infer there is recurrent innovation of selfish SNBP activities by X chromosomes, consistent with the notion of sex chromosome meiotic drive that requires silencing by endogenous RNAi. Protamines are also functionally relevant to activity of *Segregation Distorter*, an autosomal meiotic drive system in *D. melanogaster* (Gingell and McLean, 2020; Herbette et al., 2021). Moreover, a preprint by the Malik group describes further evolutionary dynamics of testis HMG box loci across the Drosophilid genus, supporting the notion that their rapid evolution is due to recurrent intragenomic conflict between sex chromosomes (Chang and Malik, 2022). We predict that hpRNAs may be employed to silence other protamine-based meiotic drive phenomenon in other species.

Within the Dox superfamily system itself, we document the innovation of novel hpRNAs that bear chimeric domain structures characteristic of coding Dox genes, but that lack the HMG box. In fact, these constitute novel hpRNAs that appear to have captured other functionalities relevant to meiotic drive (e.g. piRNA factors and TEs). Of course, TEs are intrinsically selfish elements that are targeted by host genomic defenses, most famously by piRNAs. However, recent studies provide analogous conceptual involvement for co-option of the piRNA pathway by drive systems. For example, the Paris *sex ratio* meiotic drive system utilizes a *de novo* copy of the HP1-like factor Rhino (Helleu *et al*., 2016), a central nuclear piRNA factor that defines piRNA cluster transcription (Klattenhoff *et al*., 2009; Mohn *et al*., 2014). As another example, telomeric TART elements were found to have captured a fragment of Nxf2 (Ellison et al., 2020), a piRNA-specific copy of the mRNA export machinery that gained activity in co-transcriptional silencing (Batki et al., 2019; Fabry et al., 2019; Murano et al., 2019; Zhao et al., 2019). We surmise that the capture of some piRNA factors by hpRNA loci may in fact reflect their selfish activities of such defense factors.

The adaptive deployment of hpRNAs in the testis in *D. simulans* highlights that some of the most important biologically overt manifestations of endo-RNAi cannot be studied in the major model system *D. melanogaster*. Looking to other Drosophilids, the recognition of rampant duplications of the RNAi effector AGO2 in various *obscura* clade species, resulting primarily in testis-restricted paralogs (Crysnanto and Obbard, 2019; Ellison and Bachtrog, 2019; Lewis et al., 2016a; Lewis et al., 2016b), provides a further hint into active genomic conflict scenarios that must be playing out in these species. Indeed, intragenomic conflicts that mediate aberrant *sex ratio* and/or sterility, specifically in male fathers, have been documented in the *obscura* clade (Noor, 1995; Phadnis and Orr, 2009). Overall, the Drosophila RNAi/hpRNA pathway provides a policing system that helps to surveil and control gene expression in the testis, and may provide molecular keys to unlock programs of meiotic drive.

## Materials and Methods

### *Drosophila* strains

Stocks bearing *D. melanogaster dcr-2* alleles *wIR; dcr-2[L811fsx]/CyO* and *wIR; dcr-2[R416X]/CyO* were obtained from Richard Carthew (Northwestern) (Lee et al., 2004). *D. simulans w[XD1]* wild-type strain was obtained from BestGene, Inc. and used as the control strain for mutant comparisons. *dcr-2* loss of function mutant with *DsRed* allele replacing the endogenous locus was made in the *w[XD1]* strain background as reported in (Lin *et al*., 2018). Similar to *dcr-2* DsRed mutant allele, we also generated *dcr-2* mutant allele replacing the endogenous locus with a mini-white^+^ marker for efficient selection of *dcr-2* trans-heterozygous mutants by crossing *dcr-2[DsRed]* and *dcr-2[white^+^]* parental files. All flies were reared on standard cornmeal molasses food. As *D. simulans* lack balancer chromosomes, and as *dcr2* homozygous mutants are also male sterile (Lin *et al*., 2018), we maintained *dcr-2[DsRed]* and *dcr-2[white^+^]* alleles by visual selection of markers every few generations to maintain the alleles.

### Testis dissection and RNA preparation

For *D. melanogaster* testis dissections *dcr-2* mutant testis were collected from *wIR; dcr-2[L811fsx]/dcr-2[R416X]* trans-heterozygotes and *wIR; dcr-2[R416X]/+* heterozygous flies were used as controls. For *D. simulans*, we collected testis from *dcr2[DsRed]*/[white^+^] trans-heterozygous mutants, and used the parental strain *w[XD1]* as control. Briefly, testis from 3 days old flies were extracted in TRIzol (Invitrogen) in batches of 10 flies at a time and the testis samples were flash frozen in liquid nitrogen. RNA was extracted from 25-50 testis per genotype.

### Small RNA and RNA-seq library preparation

RNA extraction was performed as described in (Lin *et al*., 2018), and the quality of RNA samples were assessed with the Agilent Bioanalyzer. RNA samples with RIN >6.5 were used for library preparation using the Illumina TruSeq Total RNA library Prep Kit LT. Briefly, for RNA-seq libraries we used 650 ng of total RNA, and we used the Manufacturer’s protocol except for reducing the number of PCR cycles from 15 as recommended to 8, to minimize artifacts that may arise from PCR amplification. We prepared stranded RNA-seq libraries for *D. simulans* and unstranded libraries for *D. melanogaster* as RNA samples were extracted and processed in different time points. Samples were pooled using barcoded adapters provided by the manufacturer and the paired-end sequencing was performed at New York Genome Center using PE75 in the Illumina HiSeq2500 sequencer.

We prepared small RNA libraries used ~20 μg total RNA, as previously described {Lin, 2018 #14350. To the total RNA pool, we added a set of 52 RNA spike-ins, spanning a range of concentrations (QIAseq miRNA Library Spike-In kit #800100). Briefly, small RNAs of size 18- to 29-nt-long small RNAs were purified by preparative PAGE. Next, the 3′ linker (containing four random nucleotides) was ligated overnight using T4 RNA ligase 2, truncated K227Q (NEB), after which the products were recovered by a second PAGE purification. 5′ RNA linkers with four terminal random nucleotides were then ligated to the small RNAs using T4 RNA ligase (NEB) followed by another round of PAGE purification. The cloned small RNAs were then reverse transcribed, PCR amplified and sequenced using P50 single-end sequencing on the Illumina HiSeq 2500 sequencer.

### 5’-seq and 3’-seq library preparation

To map 5’ ends, we used the parallel analysis of RNA 5′ ends from low-input RNA (nanoPARE) strategy (Schon *et al*., 2018). For *Dsim* libraries, testis was extracted from <1-week males and total RNA was extracted using TRIzol. cDNA was prepared using Smart-seq2 (Picelli et al., 2013) and tagmented using the Illumina Nextera DNA library preparation kit, purified using the Zymo 5x DNA Clean and Concentrator kit (Zymo Research), and eluted with resuspension buffer. For 5’-end enrichment PCR, the purified reaction was split and amplified either Tn5.1/TSO or Tn5.2/TSO enrichment oligonucleotide primer sets. PCR reaction products with Tn5.1/TSO enrichment oligonucleotide and Tn5.2/TSO enrichment oligonucleotide primer sets were pooled and purified using AMPureXP DNA beads. Final libraries were checked for quality on an Agilent DNA HS Bioanalyzer chip. Libraries with size ranges between 150 and 800 bp were diluted and sequenced to 10–15 million single-end 50-bp reads per sample using a custom sequencing primer (TSO_Seq) and a custom P5/P7 index primer mix on an Illumina HiSeq 2500 instrument.

To annotate 3’ transcript termini, we used the QuantSeq 3’ mRNA-seq library preparation REV kit for Illumina (Lexogen) with a starting material of 50 ng total RNA from *Dmel* and *Dsim* control and *dcr-2* mutant samples, according to manufacturer’s instructions. cDNA libraries were sequenced on Illumina HiSeq-1000 sequencer with single-end SE 50 mode.

### Genomic analyses of RNA-seq data in *Dmel* and *Dsim*

#### RNA sequencing analysis

Paired-end RNA-seq reads from wild-type and mutant *dcr-2* samples in *Dmel* and *Dsim* were mapped to dm6 (FlyBase) and *Dsim* PacBio assemblies (Chakraborty *et al*., 2021), respectively using hisat2 aligner (Kim et al., 2015; Pertea et al., 2016).The resulting alignments in SAM format was converted to BAM using SAMtools software (Li et al., 2009) for downstream analyses. Mapping quality and statistics were determined using the *bam_stat.py* script provided in the RSeQC software (Wang et al., 2012). Transcript abundance was determined using FeatureCounts software from the subread package (Liao et al., 2014), using *Dmel* gene annotations from FlyBase r6.25. For *Dsim*, we used both gene annotations from FlyBase and *de novo* transcript annotation using StringTie software (see details below) (Pertea et al., 2015). As FlyBase gene annotations for *Dsim* correspond to *Dsim* r2.02 assembly, we converted the FlyBase assembly annotations to *Dsim* PacBio coordinates using the UCSC liftover tool implemented in the KentUtils toolkit from UCSC (https://github.com/ENCODE-DCC/kentUtils). We combined FlyBase liftover and *de novo* annotations in *Dsim* to determine transcript abundance for RNA-seq analyses. The following description for differential gene expression (DFE) analysis is the same for *Dmel* and *Dsim* data. DFE comparing control and *dcr-2* mutant data was performed using the DEseq2 package in R (Love et al., 2014). Genes with low read counts and/or high variability among technical or biological replicates can lead to log fold change differences that are not representative of true differences. Therefore, to minimize variance, we used the log fold change (LFC) shrinkage implemented in the DEseq2 package using the ‘normal’ method described in (Love *et al*., 2014). For visualization of mapped reads, the BAM alignment files were converted to bigwig format using *bam2wig.py* script from RSeQC (Wang *et al*., 2012) and the bigwig tracks were visualized on the IGV genome browser (Robinson et al., 2011).

#### Small RNA sequencing analysis

Adapters were trimmed from small RNA sequences using Cutadapt software (https://github.com/marcelm/cutadapt); then the 5’ and 3’ 4-nt linkers (total 8 bp) were removed using sRNA_linker_removal.sh script described in (Vedanayagam *et al*., 2021) (https://github.com/Lai-Lab-Sloan-Kettering/Dox_evolution). The adapter and linker removed sequences were then filtered to remove < 15 nt reads. We mapped > 15 nt reads from Dmel and Dsim genotypes to dm6 reference genome assembly and Dsim PacBio assembly, respectively, with Bowtie (Langmead et al., 2009) using the following mapping options: bowtie -q -p 4 -v 3 -k 20 -- best -strata. The resulting BAM alignments from bowtie mapping were converted to bigwig for visualization using *bam2wig.py* script from the RSeQC software (Wang *et al*., 2012). During the BAM to bigwig conversion step, the small RNA mapping data was normalized to 52 spike-in sequences from the library (QIAseq miRNA Library Spike-In kit).

### *De novo* annotation of testis transcriptome

In addition to previously annotated transcripts/genes from the FlyBase annotation, we performed *de novo* annotation of our transcriptome data to identify additional, novel testis-expressed transcripts in *D. melanogaster* and *D. simulans*. The novel annotated transcripts were then supplemented with known annotations to make a combined set of 17285 transcripts in *D. melanogaster* and 15119 transcripts in *D. simulans*. We employed two independent, genome assembly guided transcript prediction algorithms, Cufflinks (Trapnell et al., 2012) and StringTie (Pertea *et al*., 2015). For both methods, *de novo* transcripts were predicted for each RNA-seq dataset, and a merged transcript model was generated encompassing the transcriptome from WT and mutant datasets. hpRNAs were predicted using the scheme shown in Supplementary Figure 2, and visualized using the Integrated Genomics Viewer (IGV) (Thorvaldsdottir et al., 2013). The termini of primary hpRNA transcripts were refined using the 5’-seq and 3’-seq data.

## Acknowledgements

We are grateful to members of the *simulans* clade PacBio sequencing consortium (J. J. Emerson, Amanda Larracuente, Colin Meiklejohn and Kristi Montooth) for access to *D. simulans* PacBio data at the unpublished stage. We thank Richard Carthew (Northwestern University) and the San Diego Drosophila Stock Center for fly stocks. JV was supported by a Pathway to Independence Award (NIH-K99GM137077), and work in ECL’s group was supported by the National Institutes of Health (R01-HD108914 and R01-GM083300), BSF-2015398, and MSK Core Grant P30-CA008748.

## Supplementary Figures

**Supplementary Figure 1:**
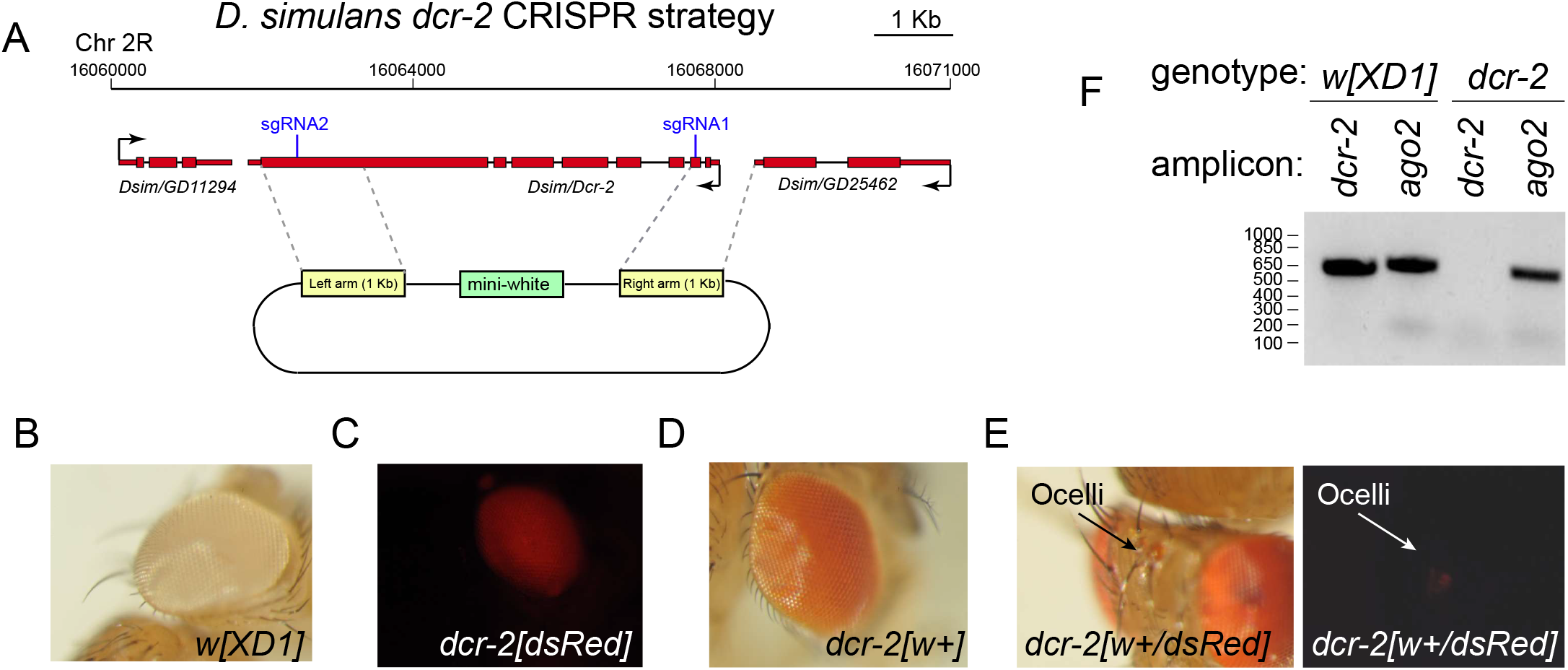
*D. simulans dcr-2* CRISPR mutagenesis strategy and validation. (A) *dcr-2* location, sgRNAs, homology donor arms with *mini-white* marker used for mutant selection. (B) *D. sim w[XD1]*, a *white* mutant background used for CRISPR. (C) *dcr-2* mutant marked by *3xP3-dsRed*; it is difficult to identify homozygous flies unambiguously. (D) *dcr-2* mutant marked with *mini-white*. (E) dcr-2 transheterozygote mutant bearing *mini-white* and *3xP3-dsRed* alleles; DsRed eyes are not very visible in a w+ background, but can be identified in the ocelli. (F) PCR genotyping and validation of *dcr-2* CRISPR mutant.

**Supplementary Figure 2:**
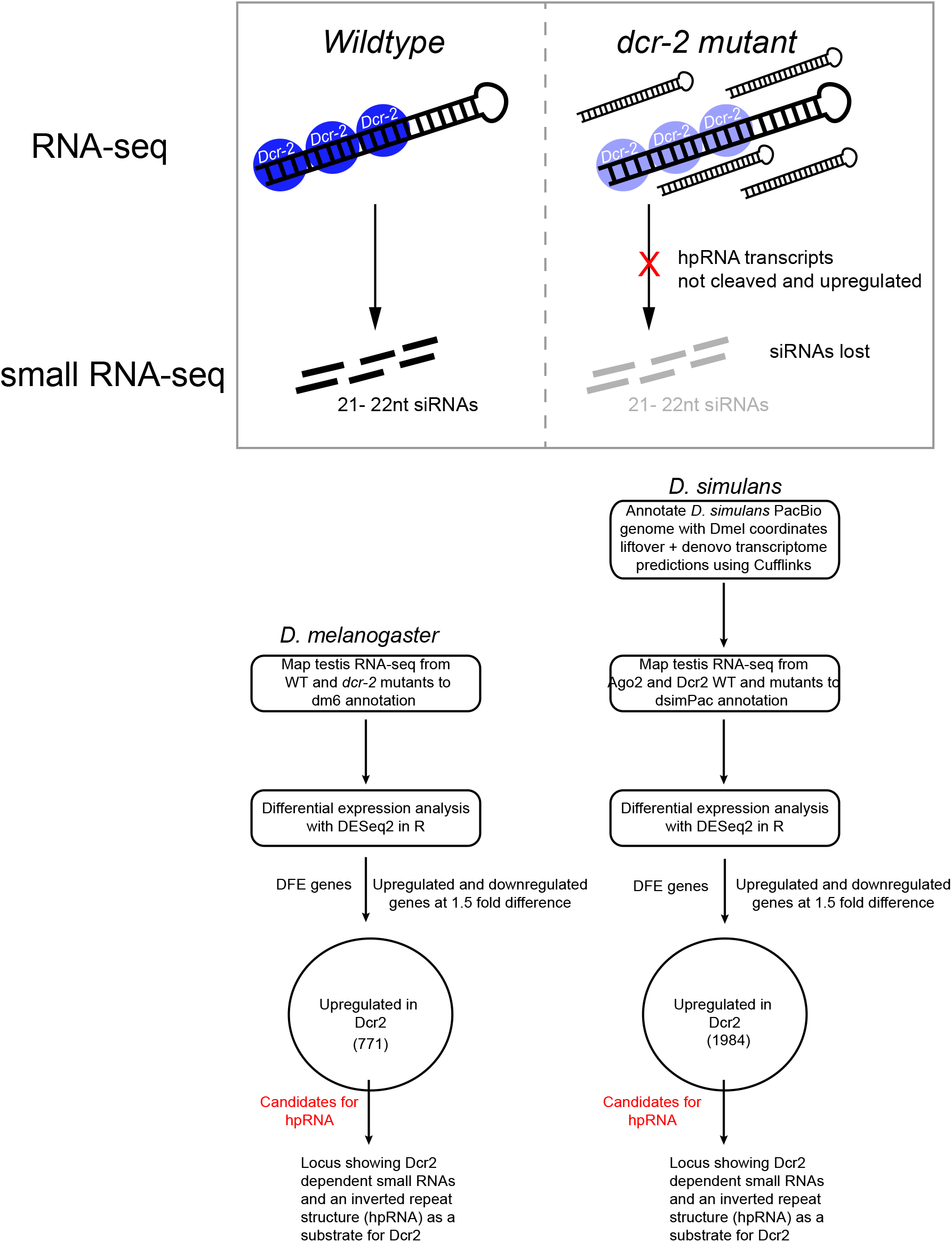
Rationale and strategy to annotate hpRNAs. (Top) Expectations of reciprocal behavior of hpRNA transcripts in wildtype and *dcr-2* mutants, with respect to RNA-seq and small RNA data. (Bottom) Overall strategy for identification of *Dmel* and *Dsim* hpRNAs. The overall procedures are similar, except that *de novo* transcriptome was generated for *Dsim*, owing to its less well-annotated genome.

**Supplementary Figure 3:**
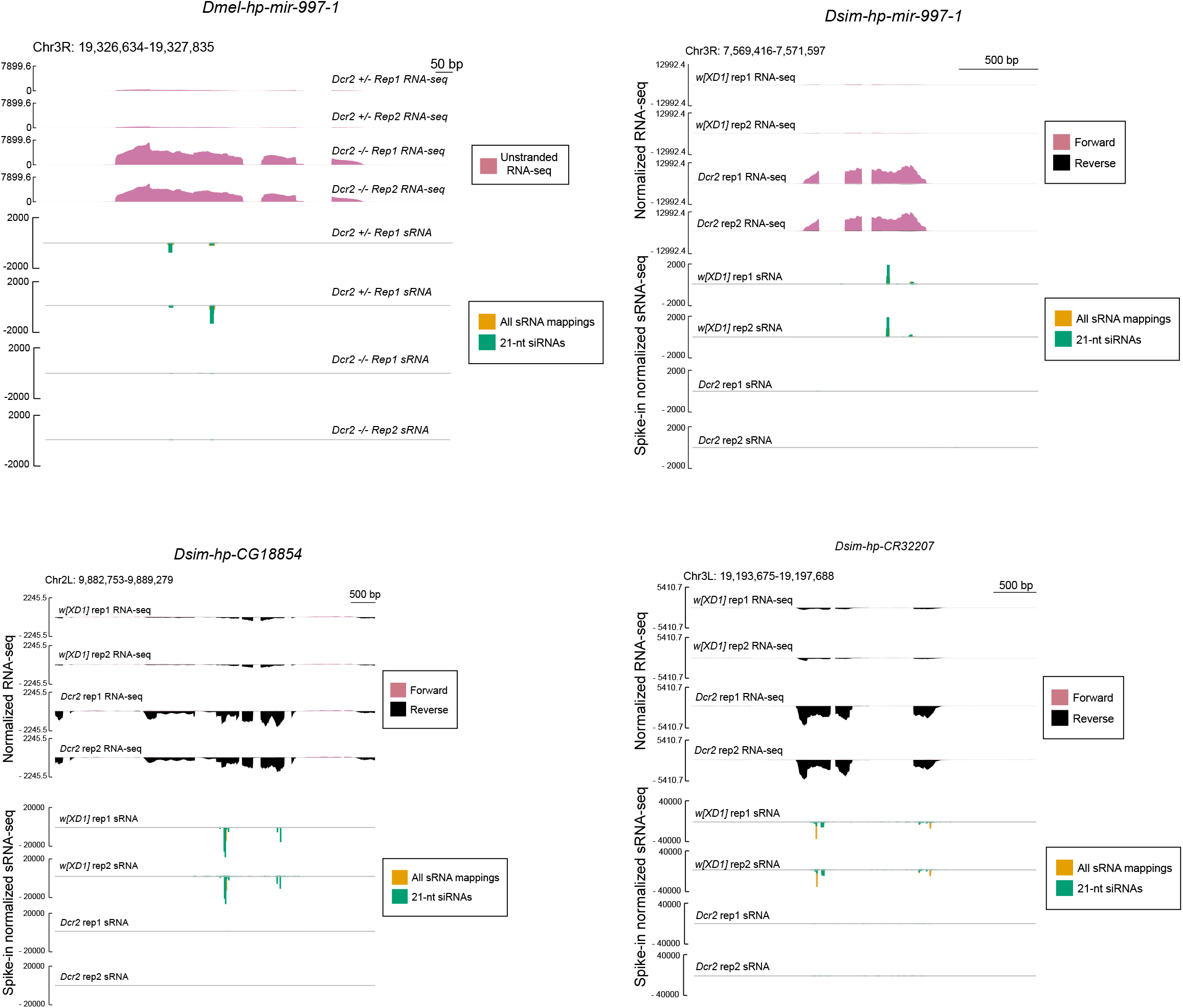
Additional examples of *Dmel*-*Dsim* conserved hpRNAs. Shown are genome browser tracks that illustrate the reciprocal behavior of hpRNAs in control vs. *dcr-2* RNA-seq and small RNA data; all of these loci are shared in the syntenic locations between *Dmel* and *Dsim*. In all cases, *dcr-2* mutants stabilize a primary hpRNA transcript while losing the ~21-nt small RNAs from the duplex regions of the hpRNA.

**Supplementary Figure 4:**
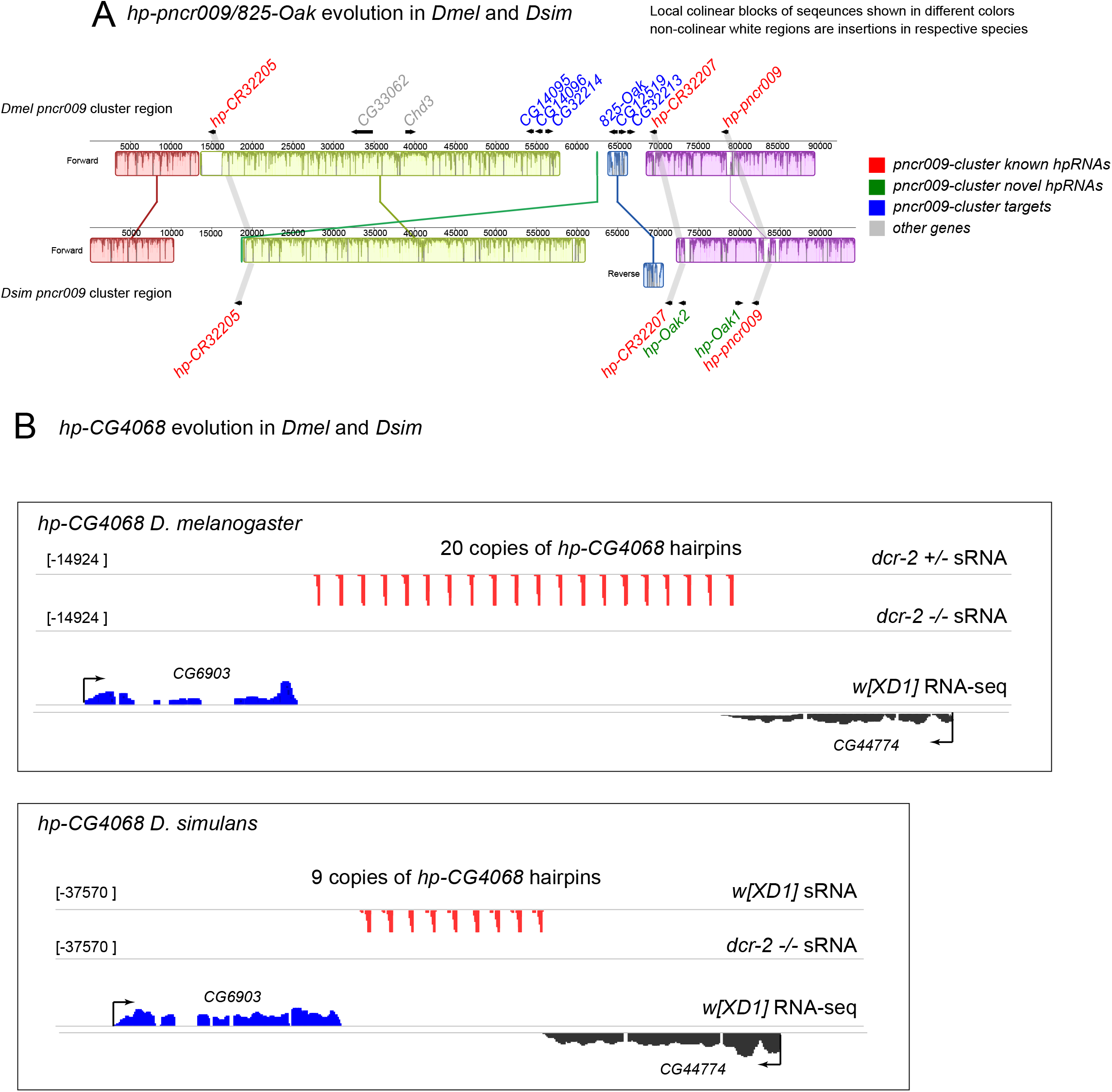
Copy number changes in hpRNA clusters in *Dmel* and *Dsim*.

(A) Synteny alignment of pncr009 hpRNA cluster region on chr3L in *Dmel* and *Dsim*. In *Dmel*, there are three hpRNAs in this region (*hp-CR32205*, *hp-CR32207*, and *hp-pncr009*, shown in red). The targets of pncr009 hpRNAs are in the vicinity of hpRNAs, and are shown in blue. Unrelated genes in the 92-kb window are shown in grey. Synteny representation is shown as local colinear blocks of sequences derived from Mauve alignment using the Geneious software. Non-colinear regions in the syntenic alignment (white regions) are insertions in respective species. Two new hpRNA paralogs in the *Dsim* pncr009 region (*hp-Oak1* and *hp-Oak2*) are shown in green. (B) IGV screenshot of hp-CG4068 cluster in *Dmel* and *Dsim*. In *Dmel*, there are 20 tandem copies of the hpRNA, while in *Dsim* there are only 9 tandem copies. Small RNA tracks show loss of siRNAs in *dcr-2* mutant testes in both *Dmel* and *Dsim*. Expression of flanking genes is shown in the RNA-seq tracks.

**Supplementary Figure 5.**
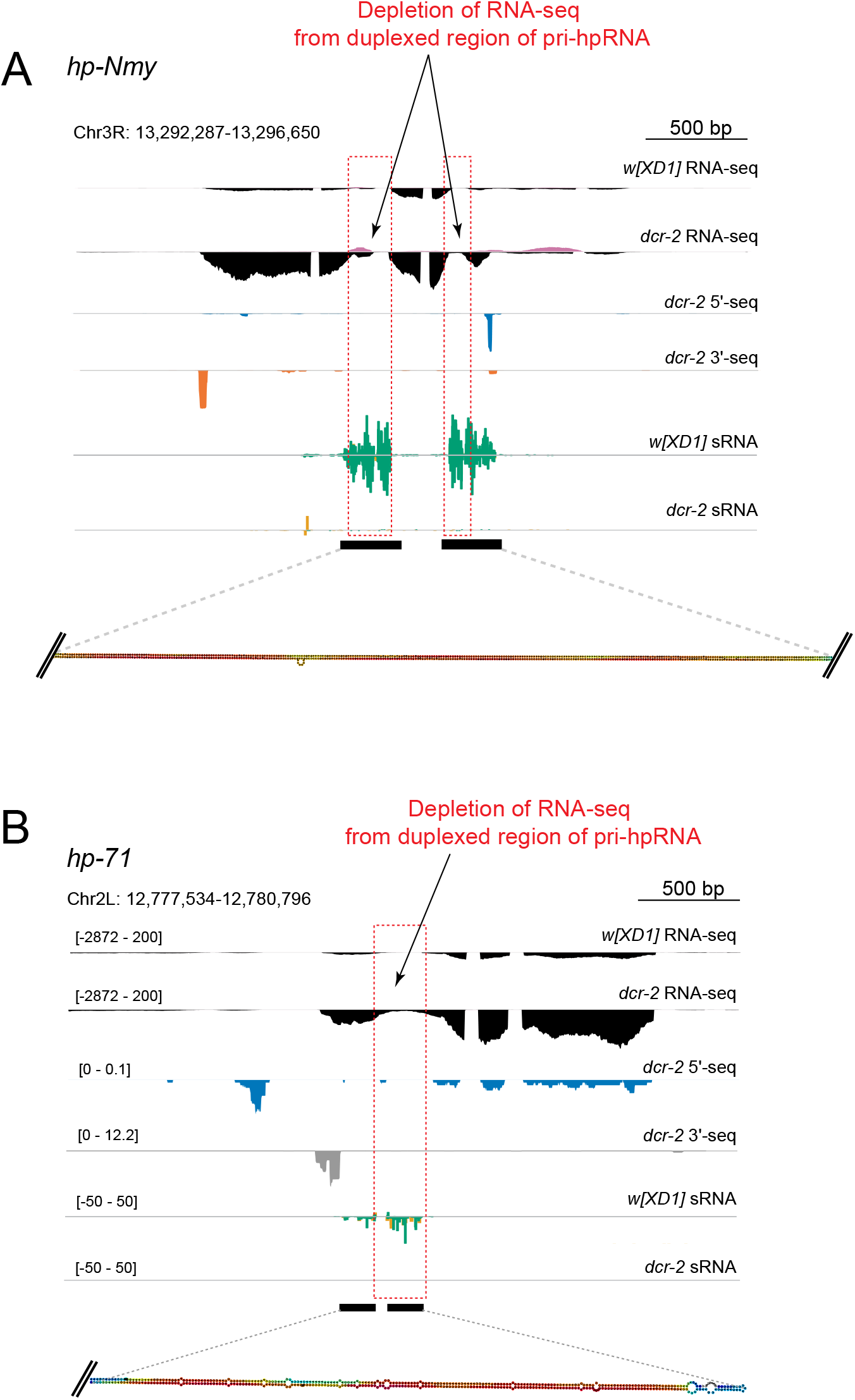
Depletion of RNA-seq data within highly duplexed regions of primary hpRNA transcripts. (A) *hp-Nmy*. The RNA-seq tracks show upregulation of pri-hpRNA in *dcr-2* mutant testis. However, the RNA-seq signal is not uniform across the primary transcript. Depletion of RNA-seq is evident within the red dotted box, corresponding to inverted repeat arms of the hpRNA. Small RNA tracks show that Dcr-2-dependent siRNAs are generated from the duplex region. (B) *hp-71* shows similar depletion of RNA-seq in the inverted repeat arm region (red dotted box).

**Supplementary Figure 6.**
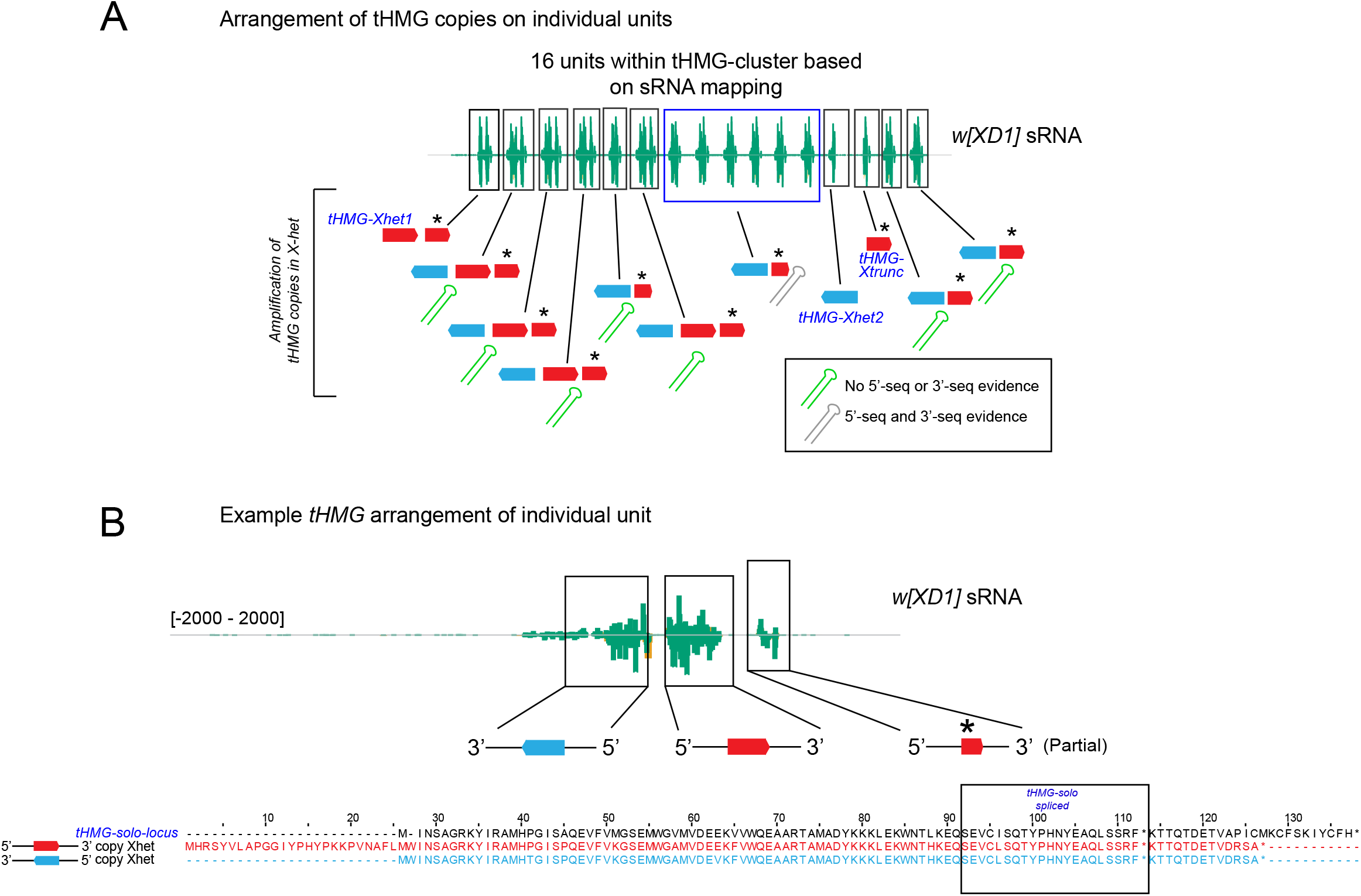
Amplification and sequence arrangement of tHMG copies in the tHMG cluster region. (A) 16 units within the X-linked tHMG cluster region are evident from small RNA mapping. Boxed regions show individual units and the tHMG-Xhet copies within each unit is shown in red and blue based on the duplicate copy orientation. Copies with an asterisk indicate partial/truncated copies of tHMG-Xhet. Within this cluster, tHMG-Xhet amplifications have birthed 13 hpRNAs and shown in green are hpRNAs for which there is absence of 5’-seq and 3’-seq evidence and 6 hpRNAs (blue box) shown in grey have 5’-seq and 3’-seq data. Shown in blue text are two copies of tHMG-Xhet (1 and 2) that are not part of an hpRNA arrangement. (B) Example tHMG arrangement of individual hpRNA unit. Shown in blue and red are orientation of paralog arrangement. Alignment of tHMG-Xhet paralogs with respect to tHMG-solo-locus is shown below. Note there is also small RNA mapping to the partial/truncated copy of the tHMG-Xhet paralog. Shown in black box on the alignment is a sequence window which is spliced at the tHMG-solo-locus.

**Supplementary Figure 7.**
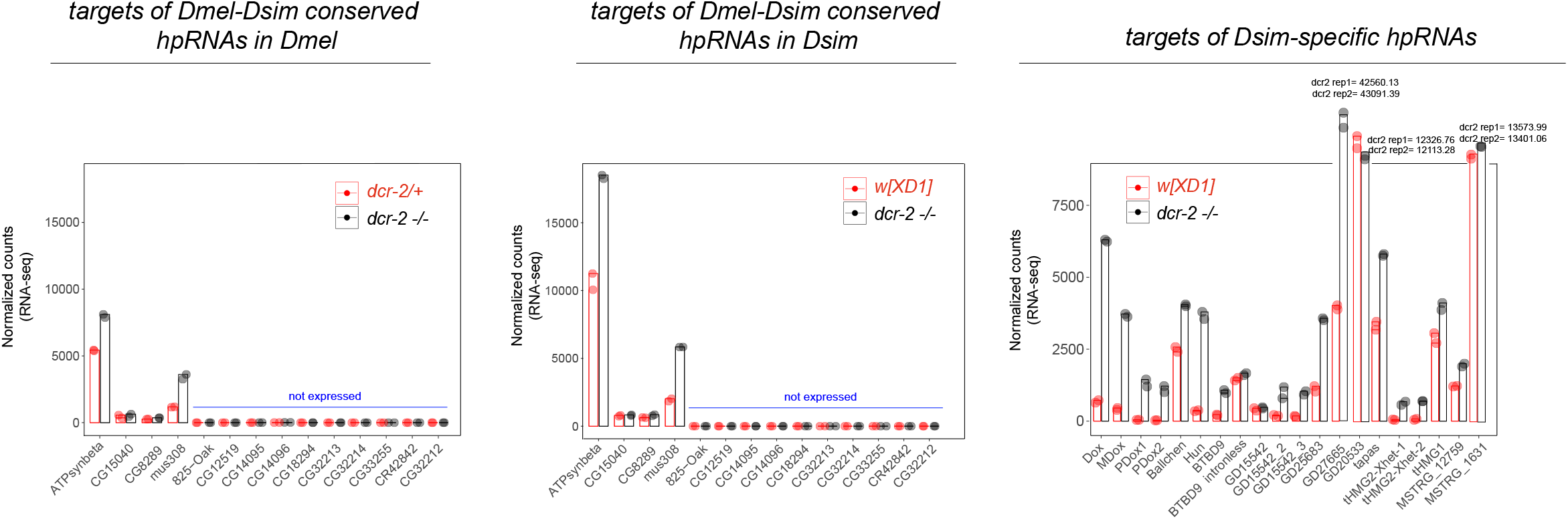
Expression of targets of *Dmel*-*Dsim* conserved hpRNAs and targets of novel *Dsim*-specific hpRNAs. Amongst conserved targets, we note that all members of the 825-Oak family show low to no expression in adult testis. Data from biologically independent RNA-seq experiments are shown as individual dots (wildtype in red and *dcr-2* mutant in black). Compared to targets of conserved hpRNAs, targets of *Dsim*-specific hpRNAs show much greater derepression in *dcr-2* mutant testis.

**Supplementary Figure 8.**
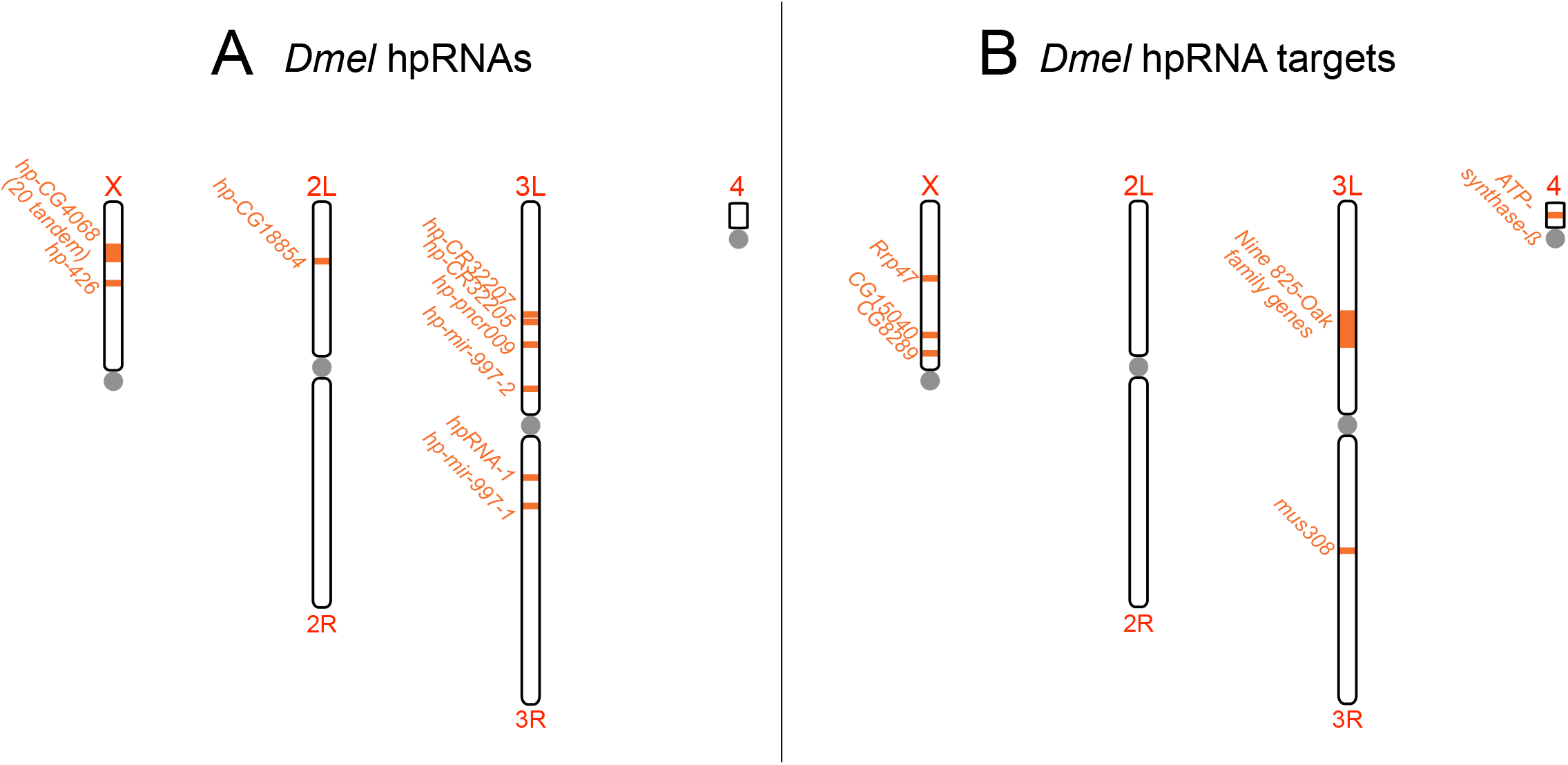
Chromosomal maps of *Dmel* hpRNAs and targets. (Left) Locations of *Dmel* hpRNAs. (Right) Locations of *Dmel* hpRNA targets. Note that all *Dmel*-specific hpRNAs have homologs in *Dsim*, whereas there are numerous *Dsim*-specific hpRNAs that are lacking in *Dmel* (see main Figures 2, 6).

